# Acute electrical neuromodulation paradigms restore firing behavior and inhibition in secondary brain injury: A computational study

**DOI:** 10.64898/2026.01.07.698101

**Authors:** Ananya Muguli, Anton Banta, Gavin W Britz, Behnaam Aazhang

**Author notes:** Corresponding author: Ananya Muguli;).

## Abstract

Secondary Brain Injury (SBI) occurs after the initial physical insult caused due to traumatic brain injuries and strokes. SBI leads to a massive loss of brain functionalities if timely intervention is not administered. This work analyses the delineated effects of three acute SBI processes in two-neuron motifs and shows that the right stimulation paradigm during specific key SBI events help restore firing rate trends in neurons. The Hodgkin-Huxley neuron is extended to theoretically model two categories of two-neuron motifs: a *feedforward excitatory motif (Motif 1: Two pyramidal neurons with glutamatergic synapses)* and a *feedforward inhibitory motif (Motif 2: One pyramidal neuron and one interneuron with GABAergic synapses)*. Three important SBI processes in the motifs were modeled: *glutamate excitotoxicity, increased extracellular potassium ion concentration*, and *cellular energy deficit*. Firing rate trend analysis is performed for increasing severity of SBI processes, and interesting points *(key events)* are identified. A wide range of ACS and DBS stimulation parameters are applied to the motifs during these *key events* to assess the firing rate response during stimulation. Increasing SBI severity caused an overall increase in excitation, synchronizing neuronal activities in Motif 1 and reducing inhibition in Motif 2. Alternate current stimulation paradigms were found to desynchronize and regulate neuronal firing in Motif 1. Deep brain stimulation parameters were found to increase inhibition in Motif 2, thereby helping to maintain the excitation-inhibition balance. The right stimulation paradigm administered at appropriate key events helps regulate neuronal firing, thereby reducing the metabolic burden on the neurons during acute SBI.

**Author Summary:** Traumatic Brain Injuries (TBIs) and strokes affect millions causing issues such as motor and speech disorders, memory and cognitive decline adding to high global economic burden. Secondary Brain Injury (SBI) is the aftermath of TBIs and strokes, which causes loss of brain functionalities, leading to disabilities. Early intervention reduces disabilities and improves quality of survivor’s life. Therapeutic electrical brain stimulation has gained prominence to help restore brain functionalities post brain injuries. It is usually administered chronically post injury, when disabilities have set. Here, we investigate if electrical brain stimulation during acute SBI leads to prevention of neurodegeneration, thereby retention of brain functionalities using biophysics. We model fundamental two neuron motifs exhibiting feedforward excitatory and feedforward inhibitory behavior (form the backbone of neuronal networks) and add SBI pathways and brain stimulation models to this scenario. On analyzing neuronal firing during various stimulation strategies at *key SBI events*, we found that specific Alternate Current Stimulation parameters can desynchronize and regulate firing, and Deep Brain Stimulation parameters could restore inhibition when the underlying neuronal properties are known and leveraged. Hence, we show that acute electrical stimulation regulates neuronal firing with appropriate stimulation parameters and SBI conditions, thereby suggesting possible effective early intervention strategies.

## 1. Introduction

Traumatic Brain Injuries (TBIs) and strokes (Ischemic, Hemorrhagic) cause the physical damage of the neural tissue, stimulating the neurodegeneration of the cells, leading to long-term issues such as motor disorders, memory and cognitive decline. TBIs are commonly caused by a violent impact to the head, thereby physically damaging the neural tissue. In 2021, it was identified that there were 37.93 million prevalent cases of TBI in the world (1), with survivors facing the aftermath of loss of cognitive capabilities and motor disorders. Strokes, on the other hand, are brain injuries caused by either blockage (Ischemic) or rupture (Hemorrhagic) of cerebral blood vessels, leading to impaired blood supply to brain parenchyma, triggering cellular death. It has been shown that each year, approximately 12 million people suffer a stroke, with nearly 94 million people being current survivors of stroke around the world. The global Disability Adjusted Life Years (DALYs) amongst the survivors is estimated to be 150 million years, leading to a high global cost of stroke of over US$890 billion (2). Hence, early intervention post brain injuries of such nature is crucial to reduce the extent of disability amongst survivors and thereby improving the quality of life.

The initial injury to the brain during TBIs and strokes is called the Primary Brain Injury (PBI) which constitutes the physical damage to neural tissue (during TBIs) or the direct damage caused due to cerebrovascular breakdown (during strokes). Secondary Brain Injury (SBI) is the indirect, delayed consequence of PBI, triggered by a cascade of pathophysiological phenomena such as neuroinflammation, oxidative stress, mitochondrial dysfunction, glutamate excitotoxicity, neurochemical imbalance, loss of ion gradients etc.(3–6), worsening the condition of the neural tissue. Neurodegeneration of large groups of cells during SBI causes loss of brain functionality. The extent of damage caused by SBI significantly determines long-term prognosis of the patients (7) and thereby DALYs.

Though SBIs exacerbate the effects of PBI, they are preventable and reversible *to an extent* with timely medical intervention (8–10). Pharmacological interventions in the past have had mixed outcomes, with only a few directed medications showing some reduction in the rate of poor outcome amongst survivors with Sub-Arachnoid Hemorrhagic stroke (SAH), while no convincing proof was found in the cases of TBIs, ischemic and Intracerebral Hemorrhagic (ICH) strokes (11). Past experimental studies in brain stimulation techniques (which involve administration of electric currents, magnetic fields, ultrasound waves to modulate neuronal firing) like Deep Brain Stimulation (DBS) of thalamus have shown to improve patient outcome in the chronic phase (12) post TBI. Administration of transcranial Direct Current Stimulation (tDCS) during sub-acute phase post TBI showed noticeable improvements in patients (13). Many protocols of Transcranial Magnetic Stimulation (TMS), and Vagus Nerve Stimulation (VNS) have shown to improve the aftermath of TBI (14). Non-invasive brain stimulation therapies like tDCS, TMS, iTBS (intermittent Theta Burst Stimulations) during chronic post stroke, when combined with physical therapy have shown improvement in motor and cognitive functions (15,16). Further, repetitive TMS (rTMS) has shown to be beneficial for motor and cognitive outcomes after stroke (17). DBS in cerebellar regions along with physical rehabilitation was found to be safe and feasible with noticeable improvement in the modulation of chronic neuroplasticity post stroke to improve upper-extremity functionalities (18). VNS is routinely used to assist post stroke recovery in patients (19) and has recently shown benefits in aggressive strokes like ICH.

Computational modeling is an effective way to understand neurodegenerative neurological conditions. The Hodgkin Huxley (HH) model (20) of a single neuron has been adapted as the base for numerous such scenarios where various pathological conditions are incorporated. The HH model was extended to capture ion dynamics during seizures, spreading depressions, increased metabolic load during SBI, and motor neuron degeneration (21–24). Stimulation techniques like tDCS, TMS (25), DBS, Temporal Interference stimulation (26,27) have been modeled in the past to understand its effects under normal and diseased conditions. The mode of mechanism of DBS in Parkinson’s was investigated using non-linear system dynamics (28). In another study (29), a two variable neuron model, Izhikevich model (30) was used to build a network of neurons to study the effect of DBS in Parkinson’s. The effects of electrical stimulation in trigeminal neuralgia network was also studied by extending the Hodgkin Huxley model (31). Overall, past works have shown that it is possible to understand neurological disorders and the effects of neurostimulation through computational modeling.

This work aims to understand the impact and mechanisms of acute electrical stimulation paradigms during SBI in two-neuron motifs, showing plausible regularization of neuronal firing behavior. Recent experimental work resonates with our work by showing that acute electrical stimulation post ischemic stroke regulates the depolarizations of neurons thereby offering a safe and effective early intervention strategy (32). Here, two electrical stimulation paradigms were modeled, namely: *Alternate Current Stimulation (ACS)* and *monophasic Deep Brain Stimulation (DBS).* The simulations show that ACS contributed to desynchronization of pyramidal neurons while DBS reduced excessive depolarizations by increasing inhibition. Two neuron motifs have not been studied in the context of SBI as per our knowledge. Understanding complex detrimental phenomena like SBI and mode of action of electrical stimulation at two neuron level would enhance our understanding of efficacious acute brain injury treatment.

## 2. Methods

### 2.1 The two-neuron motifs

Two neuron motifs (33) have been implemented as they represent the fundamental units of a network of neurons in the brain. Looking into the effects of diseases, neurological conditions, and therapeutic solutions in simple motifs will help us build intuition about the plausible underlying mechanisms at play, which aggregate and exhibit complex phenomena at the neuron population level. Motif 1 represents the *feedforward excitatory system* comprised of two pyramidal neurons interacting with an *excitatory glutamatergic synapse,* as shown in Fig 1A. The presynaptic pyramidal neuron is excited by an external source, which in turn elicits activity in the postsynaptic pyramidal neuron through glutamate release and a fixed synaptic weight. Motif 2 represents the *feedforward inhibitory system* comprised of an excitatory pyramidal neuron and an inhibitory parvalbumin interneuron interacting with an *inhibitory GABAergic synapse,* as shown in Fig 1C. Here, both the inhibitory and excitatory neurons are assumed to be excited by the same external source. The activity of the inhibitory neuron modulates the firing of the pyramidal neuron through GABA (Gamma-Aminobutyric Acid) neurotransmitter release and a fixed synaptic weight.

**Fig 1:**
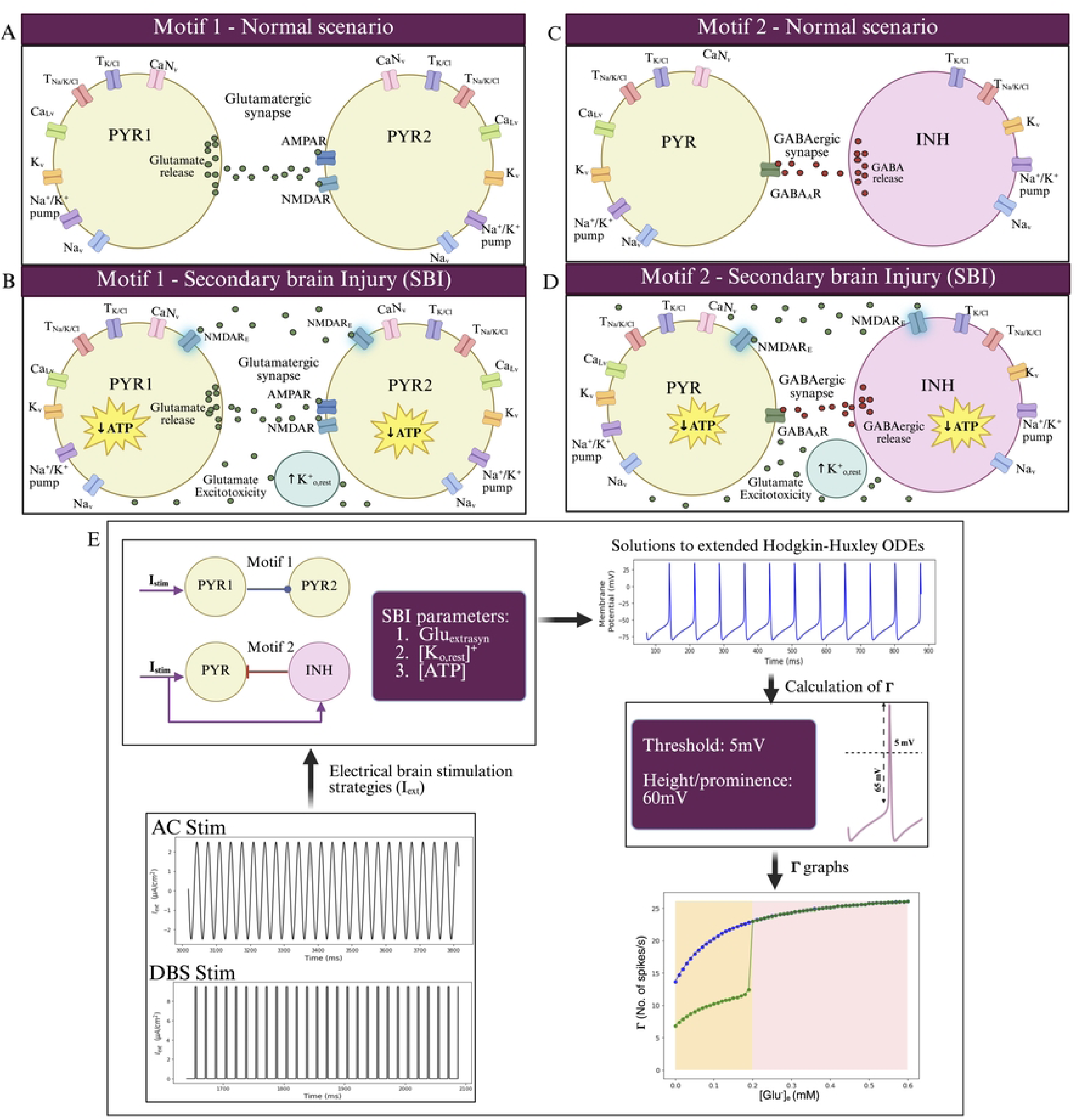
Two neuron motifs with SBI and overall research methodology. **A** Motif 1-Feedforward excitation. The two pyramidal neurons are identical, with identical ion channel behavior. The presynaptic pyramidal neuron (PYR1) elicits and modulates the activity of the postsynaptic pyramidal neuron (PYR2) through the release of glutamate neurotransmitter at the synapse. **C,** Motif 2 -Feedforward inhibition. The pyramidal neuron (PYR) is identical to the pyramidal neuron in Motif 1. The inhibitory neuron (INH) release GABA neurotransmitter at the synapse which regulates PYR firing. **B, D,** SBI pathways namely: glutamate excitotoxicity, increased *[K^+^]_o_* and ATP deficit are added to Motif 1 and Motif 2 models. **E,** The panel showcases the workflow of the analysis presented in the paper. The motifs 1 and 2 are modeled under various phases of SBI. The firing rate (Γ) is calculated from the action potential traces for a height of 5mV and a minimum prominence of 60mV. Firing rate (Γ) graphs are constructed accordingly. ACS and DBS paradigms are applied to the models during various phases of the SBI to observe the effect of electrical stimulation on firing rate and pattern of the motifs.

The *firing rate trends and patterns* of the neuron motifs are studied under the effects of the various phases of three principal SBI pathways, namely: *glutamate excitotoxicity, increased extracellular potassium concentration, and cellular (Adenosine Triphosphate (ATP)) energy deficit* (Fig 1B, D). ACS and Monophasic DBS is applied during various phases of SBI to observe the changes in the firing behavior of the motifs across various stimulation parameters. Firing rate graphs are constructed for different SBI parameter values and stimulation parameters to qualitatively analyze the effects of SBI and stimulation. Fig 1E shows the general workflow.

### 2.2 Hodgkin Huxley model of the motifs

Motif 1 and Motif 2 have been implemented by extending the Hodgkin Huxley (HH) model(20) to emulate the ionic complexity and synaptic interactions as in cortical motifs. Both categories of neurons are modeled as single compartment, point process neurons, capturing *only the temporal dynamics*. The pyramidal neuron model (34) implementation is a single compartment representing the soma. The model for perisomatic inhibitory parvalbumin interneuron is adapted from (35).

In Motif 1, the membrane potential dynamics are as follows:

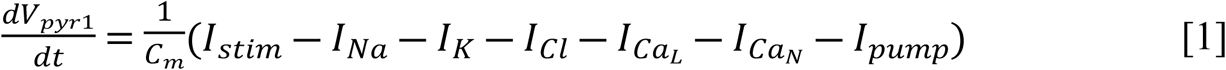

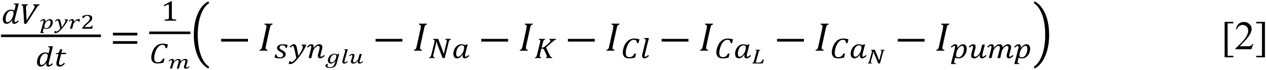

Where,

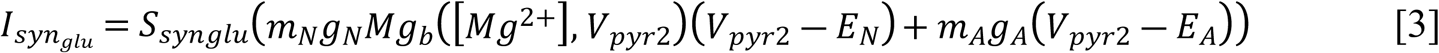

Equations [1] and [2] represent the membrane potential dynamics of presynaptic pyramidal neuron, PYR1 and postsynaptic pyramidal neuron, PYR2 neurons of Motif 1, generating two traces of action potentials. *I_stim_* is a constant input stimulus current injected to PYR1, which can be assumed to be the net activity of the upstream neurons. 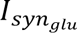 modeled in equation [3], is the synaptic current injected as input into PYR2. It is dependent on V_pyr1_ which causes glutamate release in the synapse and hence the potentiation of NMDA (N-methyl-D-aspartate) and AMPA (alpha-amino-3-hydroxy-5-methyl-4-isoxazolepropionic acid) receptors on PYR2. This causes postsynaptic membrane potential depolarization, thereby eliciting firing activity in PYR2. *E_N_* and *E_A_* are reversal potentials of NMDA and AMPA channel activity. *m_N_* and *m_A_* represent the gating variables of NMDA and AMPA receptors defined in equations [9] and [10]. *Mg_b_*([*Mg*^2+^]*V_pyr2_* term represents the role of magnesium ions as blocking agent for NMDA receptors, reducing the chances of its potentiation. This is modeled in equation [S35] (S1 Appendix).

In the Motif 2, the membrane potential dynamics are as follows:

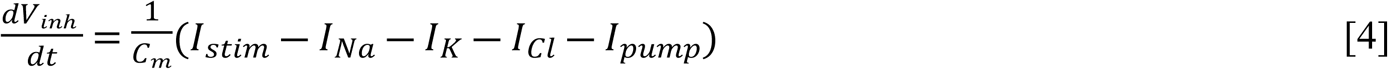

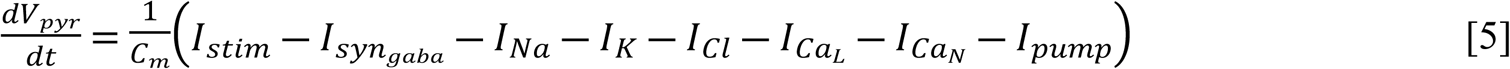

Where,

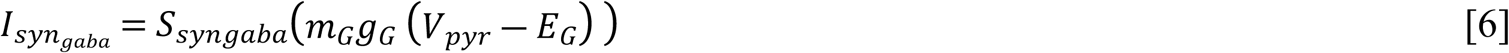

Equations [4] and [5] represent the membrane potential dynamics of INH and PYR neurons in Motif 2. *I_stim_* is the input stimuli injected to both the neurons assuming a common source. 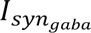 modeled in equation [6], is the synaptic current injected into PYR. The INH activity triggers the release of GABA neurotransmitter into the synapse which potentiates GABA_A_ receptors on PYR, thereby modulating PYR activity. *E_G_* is the reversal potential of GABA_A_ receptors and is dependent on the electrochemical gradient of *[Cl^-^]* across the postsynaptic membrane. Hence, *E_G_* is the same as *E_Cl_* (reversal potential of chloride ions) for the analysis in this paper. *m_G_* represents the gating variable for GABA_A_ receptor defined in equation [11].

*I_pump_* current represents the 3 Na^+^/ 2 K^+^ active pump which restores the Na^+^/K^+^ electrochemical gradient across the plasma membrane by hydrolyzing cellular energy (ATP), to transport 3 *[Na^+^]* ions out of the cell and bring 2 *[K^+^]* ions into the cell. The proper functioning of the active pump is largely dependent on availability of ATP. The equation for this current along with ATP dependency is given by :

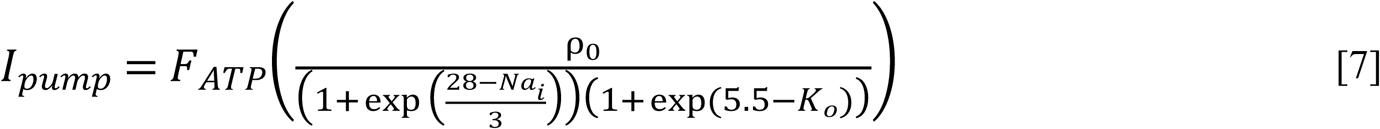

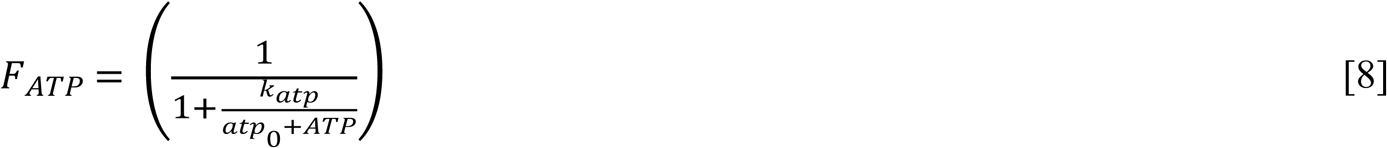

The term *F_ATP_*, is a factor that models the dependency of *I_pump_* activity on ATP concentration. The term ATP in [8] represents the current ATP concentration and *atp*_0_ represents the baseline ATP concentration. The second term represents the active pump current comprising of 3 Na^+^ ions removed from the cell and 2 K^+^ ions brought into the cell. *ρ*_0_ represents the maximum active pump current. The equations for ATP dependency and active pump current has been adapted from existing work (36).

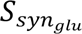 and 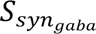 are constant parameters which indicates synaptic strength. Varying the synaptic strength will change the extent to which the presynaptic neurons influence the firing activity of post synaptic neurons in both the motifs. These parameter values are kept constants for all the analysis in the paper.

The gating dynamics of NMDA, AMPA, and GABA_A_ receptors, namely *m_N_*, *m_A_*, *m_G_* respectively were adapted from well-established kinetic models of synaptic transmission (37) as shown in Equations [9], [10] and [11]:

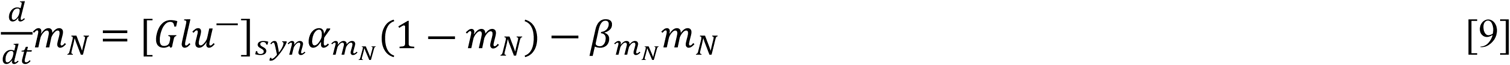

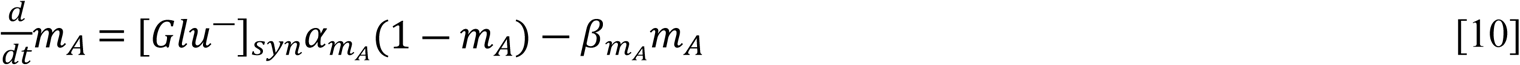

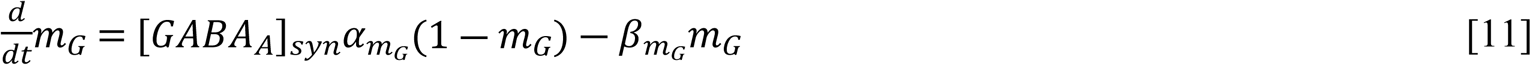

The rate constants *α*′s and *β*′s for the synaptic transmission dynamics have been adapted from standard modeling practices (37). The dynamics of the concentration of neurotransmitters released into the synapses, namely [*Glu*^−^]_*syn*_ and [*GABA_A_*]_*syn*_ is given by,

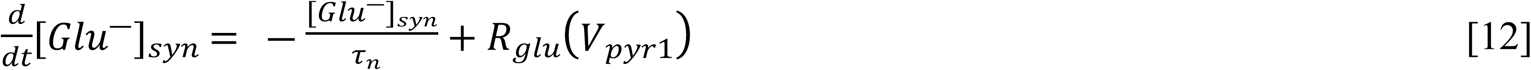

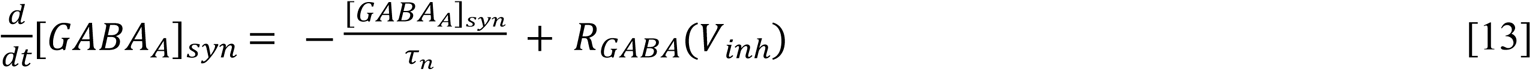

*τ_n_* is the time constant for glutamate and GABA release. The time constants have been assumed to be similar in our analysis for sake of simplicity. *R_glu_* and *R_GABA_* are the neurotransmitter release functions which are dependent on the membrane potential activity of the presynaptic neurons in both models. They are modeled for glutamate and GABA release as shown in [14] and [15],

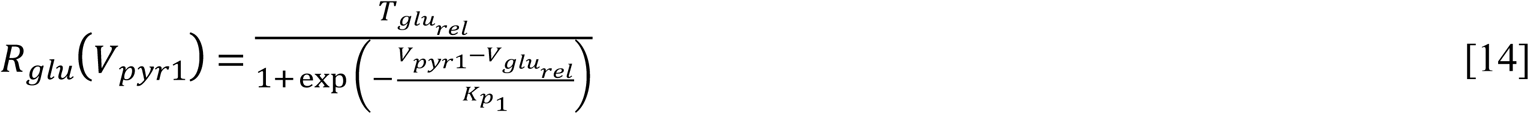

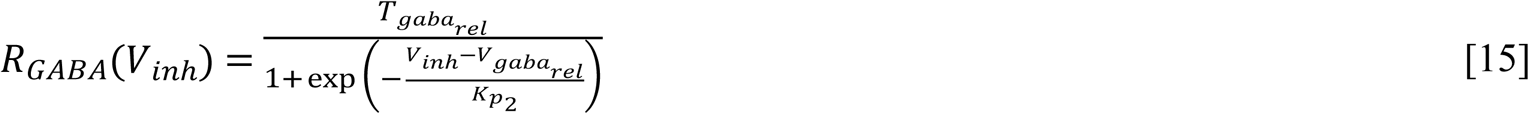

*T_glurel_* and *T_gabarel_* are constants that showcase the upper threshold of neurotransmitter release in the synapse.

*T_glurel_* and *V_gabarel_* are the presynaptic membrane thresholds at which neurotransmitter activity is elicited and is modulated by the constants *K*_*p*1_ and *K*_*p*2_.

*I_Na_ , I_K_ , I_Cl_ , I_CaL_ , I_CaN_* in Equations [1], [2], [4], and [5], are the *[Na^+^], [K^+^], [Cl^-^], [Ca^2+^]* ionic currents respectively, which are expanded in S1 Appendix. The ion concentration dynamics have been modeled to capture the effects of SBI (21,22) and electrical stimulation paradigms.

The values of constant parameters in the models have been presented in Tables S1 and S2 in S1 Appendix.

### 2.3 Modeling SBI pathways

Neurodegenerative processes such as oxidative stress, mitochondrial dysfunction, neuroinflammation, glutamate excitotoxicity, ionic imbalances set in during acute phase of SBI. Past work (22) has modeled the effects of three key SBI processes, namely glutamate excitotoxicity, increased extracellular [K^+^]_o_ and [O_2_]_o_ deficit in a single neuron model. Here, three processes were modeled, namely, glutamate excitotoxicity, increased [K^+^]_o_ and ATP deficit in two-neuron motifs to study the effects of SBI pathways and stimulation.

#### 2.3.1 Glutamate excitotoxicity

Glutamate is the most abundant excitatory amino acid neurotransmitter released by the neurons in the brain enabling synaptic transmission. Glutamate, under normal scenarios is found in high concentrations in the synaptic cleft during synaptic transmission. It triggers the synaptic NMDA and AMPA receptors in the post synaptic neurons triggering the initiation of action potentials in the post synaptic neuron. Glutamate is cleared from the synaptic cleft after the transmission by astrocytes using high glutamate affinity transporters called Excitatory Amino Acid Transporters (EAATs) (38). The hampering of glutamate release and clearance process, and breakage of blood-brain barrier (BBB) during SBIs causes the concentration of glutamate outside the synapses to increase. The extrasynaptic glutamate results in the potentiation of extrasynaptic NMDA receptors which are very permeable to *[Na^+^]* and *[Ca^2+^]* ions. This leads to excessive neuronal depolarizations and cytosolic *[Ca^2+^]* overload which further triggers other detrimental neurodegenerative mechanisms. This phenomenon is known to occur in strokes (both ischemic, and hemorrhagic), TBIs, Alzheimer’s disease, with prolonged exposures to extrasynaptic glutamate triggering cell death.

Modeling of extrasynaptic glutamate induced depolarization current 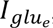, is shown in equation [17] (39). This current is added to the ODEs representing the membrane potential dynamics of all the neurons, namely *V*_*pyr*1_, *V*_*pyr*2_ in [1] and [2] and *V_pyr_*, *V_inh_* in [4] and [5]. This current is also added in equations [S19] and [S25] in S1 Appendix to account for the changes in *[Na^+^]* and *[Ca^2+^]* concentration dynamics during excitotoxic conditions.

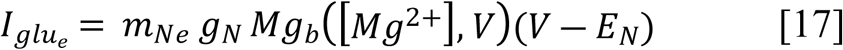

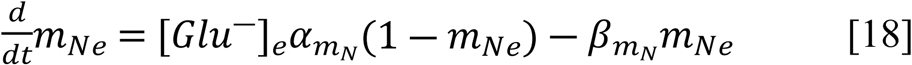

*m_Ne_* represents the gating dynamics of extrasynaptic NMDA receptors, which depends on the extrasynaptic glutamate [*Glu*^−^]*_e_* concentration. For the sake of simplicity, *α*, *β* in equation [18] is kept the same as the *α*, *β* in [9]. For INH during excitotoxicity, *g_N_* is multiplied by 0.5 to account for the smaller size of parvalbumin interneuron. [*Glu*^−^]*_e_* is the parameter used to study the effects of increased extrasynaptic glutamate concentrations by varying it across a range of values from 0mM to 0.6mM.

#### 2.3.2 Increase in extracellular potassium ion concentration ([K^+^]_o_)

During SBI, extracellular potassium ion concentration (*[K^+^]_o_*) increases due to various reasons, such as improper *[K^+^]_o_* glial uptake mechanism, neuronal hyperexcitability, and neuronal discharges. It has been shown in cases of mild TBI and concussions, massive neuronal discharges cause an increase in *[K^+^]_o_* (40,41). During ICH, where a ruptured blood vessel bleeds into the brain tissue, breakage of neuronal and glial cell membranes due to PBI induced by the hematoma buildup and BBB disruption cause a release of *[K^+^]_o_* from these cells (42), thereby worsening SBI. In ischemic strokes, where the oxygen supply to a part the brain is cutoff due to a blocked blood vessel, a rapid increase in *[K^+^]_o_* occurs due to failed ion pumps and neuronal hyperexcitability (43). Overall, increase in *[K^+^]_o_* triggers Cortical Spreading Depressions (CSD) and neuroinflammation (44). To model this phenomenon in our simulations and analysis, the parameter *[K^+^]_o,rest_* was varied across a range of values varying from 2mM to 11mM. This evidently affects the *[K^+^]_o_* dynamics modeled in equation [S21] in the S1 Appendix. Overall, such a scenario contributes to ionic imbalance and a change in the resting membrane potential of neurons.

#### 2.3.3 Cellular energy (ATP) deficit

During SBIs, mitochondrial dysfunction and cytosolic *[Ca^2+^]* overload causes significant reduction in the production of ATP in the cell which is very vital in maintaining the resting membrane potential of a neuron and the ion gradients in the cellular environment. Reduction in ATP causes failure of active pumps in both neurons and glial cells, thereby leading to ionic imbalance and chronic hyperactivity of neurons which further increases the metabolic demand in the cell, creating a detrimental positive feedback loop (24). This ultimately leads to cell death. Further, when other conditions such as glutamate excitotoxicity occur along with ATP deficit, it causes an immense metabolic load, which the cell, in most scenarios, cannot sustain under prolonged conditions of such nature. Here, we have modeled this phenomenon by varying the values of the parameter ATP from 0.01μM to 30μM, which in turn changes the value of *F_ATP_* in [8]. The analysis does not focus on the ATP production-consumption dynamics but focuses on the trends of firing behavior under scenarios of normal and abnormal ATP availabilities. The constant parameters in [8] can be modified and adapted to specific cellular energy consumption patterns as per the need.

### 2.4 Modeling electrical stimulation

Electrical stimulation paradigms have been used for therapeutic purposes for numerous neurodegenerative conditions, but the effect of stimulation parameters (frequency and amplitude of stimulation) for the given condition, situation and the patient varies widely due to numerous underlying factors. As discussed before, various past works have modeled brain stimulation paradigms (electrical (31), magnetic (25) and ultrasound (45)) and have incorporated it in the HH model as an electric current added to the membrane potential dynamics. Here, *I_ext_* represents the external electrical stimulation current. In our models, this current is added to the equations representing membrane potential dynamics: Equations [1], [2], [4], [5]. For the analysis, two types of electrical stimulation waveforms were modeled: ACS waveform and monophasic DBS waveform as shown in Fig 1E.

In equation [19], *A_ac_* represents the amplitude of AC stimulation and *f_ac_* represents the frequency of AC stimulation. For the analysis, *A_ac_* was varied in the range 0.5-8.5μA/cm^2^. The values of *f_ac_* were chosen from a discrete set of 14 frequencies: [0.5 1, 2, 5, 10, 15, 20, 25, 30, 50, 60, 100, 120, 140, 150] Hz. These frequencies were chosen based on past experimental work in AC stimulation.

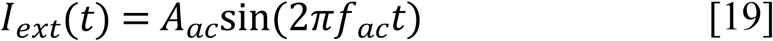

Equation [20] models monophasic DBS-like stimulation where *A_dbs_* represents DBS amplitude and *f_ac_* represents the DBS frequency. The function *heav* in [20] represents Heaviside step function. Operator *heav*, produces rectangular pulses for the width modulated positive phases of the sine function. The parameter *a* in [20] determines width of the pulses. The values for *a* varies from 0.0001 ≤ *a* ≤ 1, with higher values closer to 1 causing the pulses to be very narrow, and lower values producing broader pulses. To obtain DBS like narrow pulses, the value of *a* was fixed for all simulations at 0.96.

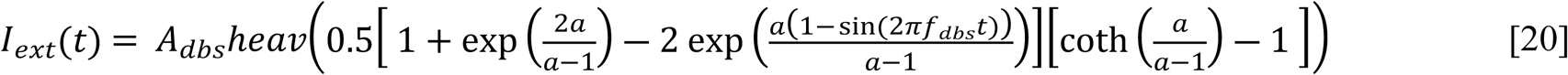

Due to the nature of [20], for DBS only two stimulation parameters can be varied, namely the amplitude, *A_dbs_* and frequency *f_dbs_*. Varying parameter *a* doesn’t lead to varying of the width of pulses tractably in the μs timescale (width of DBS pulses vary in μs range), due to which, the pulse-width parameter was eliminated for analysis. The values of *A_dbs_* vary from 0.5-8.5μA/cm^2^, matching the amplitude range used in ACS simulations. The values of *f_dbs_* were chosen from a discrete set of 7 standard frequencies that are commonly experimented during DBS procedures: [50, 60, 75, 100, 120, 140, 150] Hz.

Further, it is to be noted that the unit of *I_ext(t)_* is μA/cm^2^ because conventionally in HH models, the unit of all the currents added to the membrane potential dynamics is μA/cm^2^.

### 2.5 Simulation of motifs and calculation of firing rate

Hodgkin Huxley (HH) model is a system of Ordinary Differential Equations (ODEs), which on solving provides traces for all the variables modeled. Our extended HH model for Motif 1 comprises of two identical pyramidal neuron motifs, the equations for both the neurons being the same. These two neurons are coupled via glutamatergic synapse as discussed in the previous section. Overall, the system modeling Motif 1 consists of 22 ODEs. Similarly for Motif 2, the parvalbumin interneuron communicates with the pyramidal neuron using a GABAergic synapse. Overall, the system modeling Motif 2 consists of 19 ODEs. To solve the ODEs, a python non-linear system dynamics package called PyDSTool (46) was used. As these models were identified to be stiff systems due to varying timescales of the equations, AUTO based *Radau* (47) solver (A *Runge Kutta* based solver with adaptive time steps) was used for our simulations. The models were simulated for a total of 10000ms (10s). Due to non-stochastic nature of the systems, a duration of 10000ms (10s) was deemed sufficient to calculate a stable firing rate value for the entire duration of simulation. Solution to the ODEs gives a set of time series plots for each of the motifs, with traces of action potentials for both the neurons being one of the solutions.

Firing rate is the chosen tool for qualitative analysis because of the ease of interpretability. Further, in most scenarios, stable firing rate trend is an indicator neuronal health with dysfunctional diseased neurons deviating from such trends. Hence, by restoring the firing rate patterns of dysfunctional neurons, one could possibly reduce the chances of neurodegeneration.

The firing rate is calculated from action potential traces as shown in Fig 1E. An action potential is considered a valid spike when it fires above a height of 5mV (*V* ≥ 5*mV*) and has a minimum prominence of 60mV (|max (*V*) ― min (*V*)| ≥ 60). Overall, for a given simulation of non-stochastic ODEs, the firing rate is calculated as:

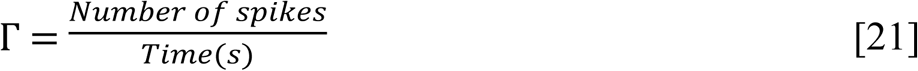

Firing of a neuron is usually considered a binary (all-or-none) phenomenon for the sake of simplicity. But neural populations under SBI is subjected to cellular energy deficit, ionic imbalances, oxidative stress, excitotoxicity and other pathways that lead to neurodegeneration. Such conditions could change the width, height and shape of the action potential. What can be considered a valid firing and calculation of Γ under such situations have not been discussed in past works. Further, such situations also change the resting membrane potential, which changes neural excitability. Hence, defining Γ calculation parameters (such as height and prominence) for such studies is subjective and should be decided after repeated evaluations of preliminary results.

## 3. Results

Understanding SBI processes independently helps in developing novel treatment methods targeting such conditions. Though brain stimulation techniques have been employed to handle the aftermath of SBIs, the mode of action and suitability of stimulation paradigms has always been debated due to varied effects which heavily depend on the state of neurodegeneration and underlying confounding factors such as genetic conditions.

Hence, the two goals of this work are:

1. *Delineate the effects* of the three SBI processes, namely: Glutamate Excitotoxicity, increased *[K^+^]_o_*, and ATP deficit by qualitatively analyzing the *Γ trends* of the processes in Motif 1 and Motif 2. Understanding these processes independently, gives us an insight into their influence on the two different neuron motifs.
2. Prove qualitatively that the *right set of DBS and ACS parameters*, at *specific phases* along acute SBI processes can help *restore Γ* in both motifs, thereby preventing irreparable damage at the cellular level.

To achieve goal 1., the *key SBI parameters* → [*Glu*^−^]*_e_*, *[K^+^]_o,rest_*, ATP were independently varied across a wide range of values to showcase each of the SBI processes. Γ of both *presynaptic and postsynaptic neurons* are calculated for each of these parameter values and trends are qualitatively analyzed for both the motifs. To achieve goal 2., at certain *key events and phase transitions* in Γ trends associated with the SBI parameters, *ACS and DBS paradigms* are independently applied for various amplitudes (*A_ac_*, *A_dbs_*) and frequencies (*f_ac_*, *f_dbs_*). This will generate a *heatmap of firing rate responses* for each of the neurons in the motifs for various stim amplitudes (y-axis) and stim frequencies (x-axis) at key events of SBI process. Γ during stimulation is compared with Γ before stimulation at the same key event to qualitatively assess the effect of stim parameters.

Based on trends observed and general response of the motifs to stimulation paradigms, Γ graphs were sectioned into different phases, namely: *Normal phase*, *Subcritical phase (Self restoring), Subcritical phase (Non self-restoring),* and *Critical phase* as seen in Fig 2. This enables us to qualitatively describe motif behavior during the progression of SBI processes and application of electrical stimulation. Further, the results are mostly discussed in terms of the *firing behavior of pyramidal neurons* in both Motif 1 and Motif 2 because they are structurally bigger, highly connected and are primarily responsible for downstream propagation of neural activity. Through goal 1., a clearer understanding of the behavior of the motifs under different values/intensities of each SBI process is provided. Through goal 2., the effects of ACS and DBS paradigms at key events along the progression of a specific SBI process is analyzed.

**Fig 2:**
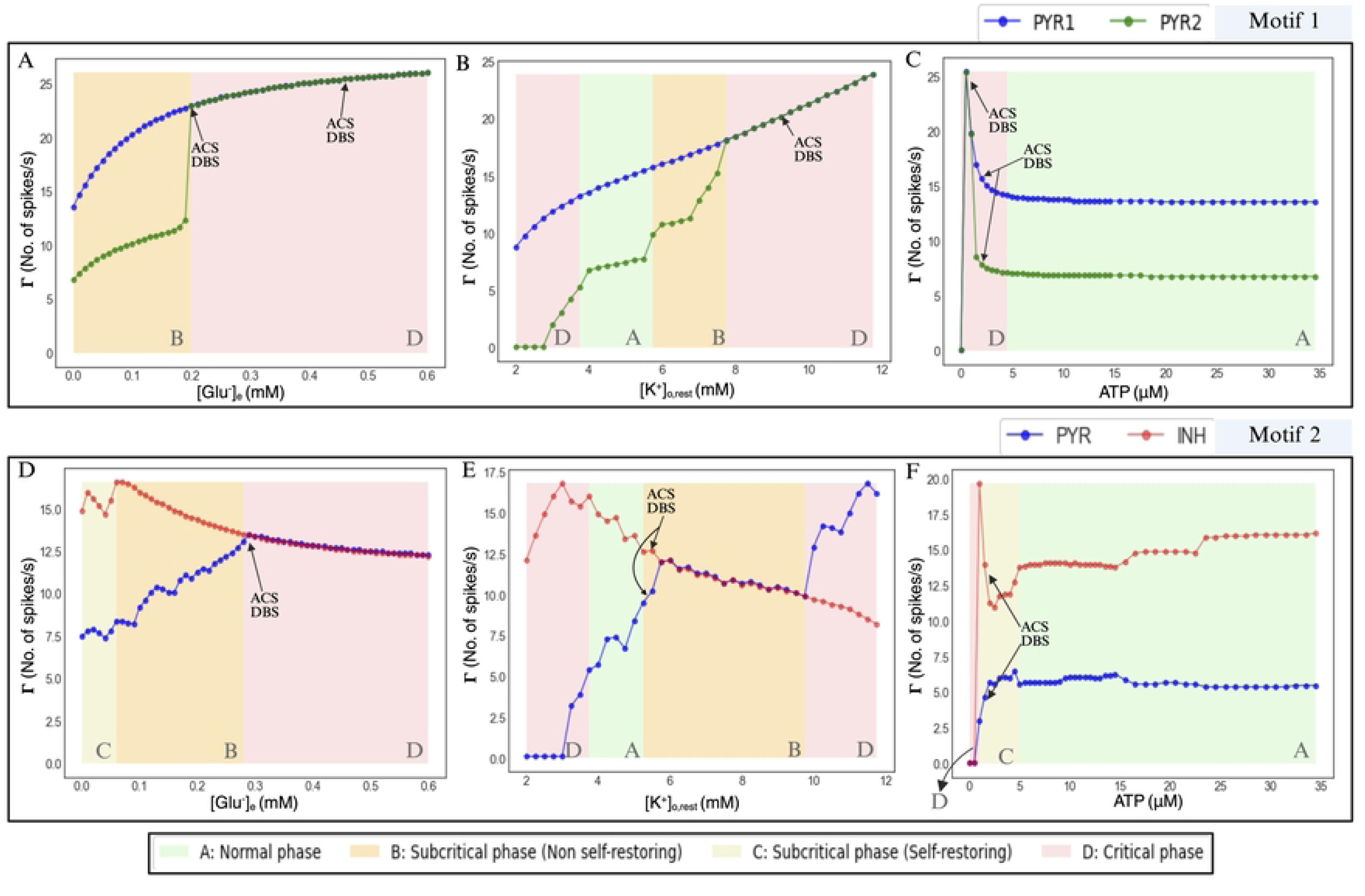
Γ trends of motifs during for various SBI processes. The *top* panel shows the Γ trends for the variation in 3 SBI parameters in Motif 1. The *bottom* panel shows the same for Motif 2. **A,** The Γ trends of PYR1 and PYR2 with varying [*Glu*^−^]*_e_*. The application of stimulation has been marked at 0.2mM and 0.45mM. **B,** The Γ trends of PYR1 and PYR2 with varying *[K^+^]_o,rest_* . The application of stimulation has been marked at 10mM. **C,** The Γ trends of PYR1 and PYR2 with varying ATP values. Application of stimulation has been marked at 0.51μM and 2.01μM respectively. **D,** The Γ trends of PYR and INH with varying [*Glu*^−^]*_e_*. Stimulation application has been marked at 0.29mM. **E,** The Γ trends of PYR and INH with varying *[K^+^]_o,rest_* . DBS and ACS application has been marked at 5.8mM. **F,** The Γ trends of PYR and INH with varying ATP values. Stimulation application has been marked at 1.51μM. In the *top* panel, the feedforward excitatory system shows increased firing in *Subcritical and Critical phases* and coherent firing of neurons mostly in *Critical Phases*. Since the system lacks an internal regulatory/self-restoring mechanism, it succumbs to excessive synchronized depolarizations due to the SBI process, unless an external intervention is administered. Both ACS and DBS paradigms were applied at *key events* to both PYR1 and PYR2 neurons. In the *bottom* panel, the PYR neuron benefits from the feedforward inhibitory system of Motif 2. The system curbs excessive depolarizations in the early stages of certain SBI processes enabling self-restoration in early stages of Subcritical phase. Latter stages of the SBI processes is marked by reduced INH activity, thereby tipping the excitation-inhibition balance towards a more excitatory behavior. ACS and DBS paradigms were applied based on the stimulation strategies discussed in Section 3: PYR-INH stim, PYR stim, and INH stim.

### 3.1 Firing rate trends during the SBI processes

#### 3.1.1 Glutamate excitotoxicity

To simulate glutamate excitotoxicity, the parameter [*Glu*^−^]*_e_* was varied in the range 0 to 0.6 mM in steps of 0.01mM and keep all other parameters constant. Under normal scenarios, [*Glu*^−^]*_e_* is 0mM. The condition of the neurons progressively worsens with increase in [*Glu*^−^]*_e_* caused due excessive depolarizations. For each [*Glu*^−^]*_e_*, the solutions to the ODEs are computed and Γ is calculated.

Fig 2A shows the effect of [*Glu*^−^]*_e_* in Motif 1. As the value of [*Glu*^−^]*_e_* increases, the overall Γ trends of both PYR1 and PYR2 increase monotonically due to increased potentiation of extrasynaptic NMDA receptors in both the neurons, which causes increased depolarizations of the neural membranes leading to hyperactivity. As the input to PYR1 is the constant *I_stim_* and the input to PYR2 is 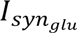 at a certain synaptic strength 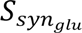, the firing rate of PYR2 is lower than the firing rate of PYR1 until the [*Glu*^−^]*_e_* value of ∼0.2mM. At [*Glu*^−^]*_e_* ≥ 0.2*mM*, the firing rate trends of the two neurons converge, exhibiting coherence in the spiking activity. It can be seen in Fig S1A (S1 Appendix), at [*Glu*^−^]*_e_* = 0mM, PYR2 fires at a lower frequency than PYR1, firing at every alternate spike of the latter. After the sharp coupling of Γ trends at ∼0.2mM, the trend saturates for [*Glu*^−^]*_e_* ≥ 0.45*mM*. Experimentally, it has been shown that the firing rates saturate at high values of [*Glu*^−^]*_e_* and sustained exposure to such concentrations causes accelerated cell death (48). For 0 ≤[*Glu*^−^]*_e_* ≤ 0.2mM, the system is in *subcritical phase (Non self-restoring)* because the increasing [*Glu*^−^]*_e_* causes increase in neuronal firings in both neurons and due to the overall excitatory nature of the system, there is no *inherent ‘inhibitory’* or a *‘regulatory’* mechanism to check the hyperactivity of the neurons. When [*Glu*^−^]*_e_* ≥ 0.2mM, the system enters *critical phase*, because the neurons are exhibiting coherent firing at high frequency (Fig S1B (S1 Appendix)), which inherently changes the nature of information propagation. The point at which neuronal firings get coupled (here, at [*Glu*^−^]*_e_* = 0.2mM) depends on 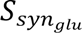 value, which though has been kept fixed for the entire analysis, was varied to test this point. Due to excessive coherent firings in such motifs, the neurons could propagate non-epileptiform discharges and succumb to high metabolic demand.

In Motif 2, for a given 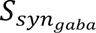, under normal conditions, Γ of PYR is lower than that of INH as seen in Fig 2D and Fig S2A (S1 Appendix). With increasing [*Glu*^−^]*_e_*, Γ of PYR neuron increases as expected but is not perfectly monotonic. In the range 0 ≤[*Glu*^−^]*_e_* ≤ 0.06mM, Γ of PYR is controlled by INH activity through the GABAergic synapse forming the *Subcritical phase (Self restoring)* as the conditions are not ideal, but the inhibitory mechanism in place counteracts the consequences of the SBI process. The phase 0.06*mM* ≤ [*Glu*^−^]*_e_* ≤ 0.29*mM*, has been marked as *Subcritical phase (non self-restoring).* Here we see a gradual lowering of INH activity and overall increase in PYR activity (49). In the implementation of INH model, the scaling factor Φ (S1 Appendix) is used to modulate the dynamics of *[Na^+^]* ion channels. Thus, a huge influx of *[Na^+^]* ions due to extrasynaptic NMDA receptor potentiation reduces the firing rate of INH. The reduced inhibition, added with high [*Glu*^−^]*_e_* conditions increases the net PYR activity, disabling self-regulation. The range [*Glu*^−^]*_e_* ≥ 0.29*mM* is the *Critical phase*, wherein the Γ trends of PYR and INH being similar, saturate (Fig S2B (S1 Appendix)) with no improvement in INH activity. The start and continuation of this phase was found to be dependent on *I_stim_*, which balances the effect of INH on PYR. Overall, Motif 1 being a wholly excitatory system is more susceptible to glutamate induced hyperactivity and further excitotoxic cell death. The PYR in Motif 2, exhibits some resilience to glutamate induced hyperactivity, but the eventual loss in inhibition, thereby the *excitation-inhibition balance*(50) causes PYR activity to increase, altering the systemic behavior.

#### 3.1.2 [K^+^]_o_ increase

To simulate the influence of extracellular potassium, *[K^+^]_o,rest_*parameter was varied from 2.0mM to 11.8mM, keeping all other parameters constant. The range 3.6mM to 5.6mM (5.2mM for Motif 2) of *[K^+^]_o,rest_* , indicates the normal range, thereby representing the *Normal phase* in both motifs.

Fig 2B shows the Γ trends in Motif 1 for variations in *[K^+^]_o,rest_*. During SBIs, the firing rate of the neurons increase (51,52) with increasing *[K^+^]_o,rest_* which has been shown in Fig 2B. PYR1 exhibits an almost monotonic linear increase in Γ, with no signs of saturation at the upper *[K^+^]_o,rest_* range. PYR2 behavior can be assumed to be a more realistic imitation of a pyramidal neuron due to being at the receiving end of a synapse. On observing the Γ trends of PYR2, the firing behavior of the motif is resilient for relatively small changes in *[K^+^]_o,rest_* in *Normal phase*. Post this, with a steep increase in firing rate, the motif enters *Subcritical phase (Non self-restoring)*. Here, the motif again maintains Γ for small changes in *[K^+^]_o,rest_*, but a further steep increase in the trends ultimately causes the merging with the Γ trend of PYR1 at 8mM, hence entering the *Critical phase*. The Fig S3A (S1 Appendix) shows the plot of action potential traces at 6mM where PYR2 is not coupled with PYR1 and shows deformities in action potential shape which occurs due to the changing ionic environment. In Fig S3B (S1 Appendix), the firings of PYR1 and PYR2 are coherent with increased Γ. In severe cases of ischemia and brain traumas, the values of *[K^+^]_o,rest_* goes well beyond 12mM, sometimes up to 30-40mM, which was not simulated due to computational constraints (53). Overall, sustained electrochemical imbalance marked by high levels of *[K^+^]_o,rest_*is detrimental to the neurons, eventually initiating cortical spreading depression. The region before the *Normal phase* has been marked as *Critical phase*, because low values of *[K^+^]_o,rest_*, is as detrimental as high values of *[K^+^]_o,rest_* and does occur in certain cases of TBIs(54). As seen in Fig 2B, this phase shows a reduction in Γ, and a near complete suppression in PYR2.

In Motif 2, under normal situations, there is a definitive balance between the excitation and inhibition in the system. But increasing *[K^+^]_o,rest_*causes an overall reduction in INH neuron activity and massive increase in the PYR activity. The *Normal phase* of the system is marked by higher INH activity modulating the activity of PYR activity. Unlike PYR neurons that exhibit some resilience in firing behavior with minor changes in *[K^+^]_o,rest_*, INH does not. Increase in *[K^+^]_o,rest_* causes reduction in INH activity, thereby increasing PYR activity. In *Subcritical phase (Non self-restoring)*, INH and PYR neurons appear synchronized, where both PYR and INH have same Γ (Fig S4A (S1 Appendix)). This effect occurs due to the value of *_stim_* chosen for our simulations, where the net excitatory input to both neurons and inhibitory effect on PYR balance out. Changing the value of *I_stim_* changes the range of *[K^+^]_o,rest_* values for which INH and PYR have the same Γ trends. Post 9.8mM, PYR neuron activity breaks off from the inhibitory effect of INH and exhibits a steep increase in firing with a steady decrease in INH activity marking the *Critical phase*. At this stage, we observe a breakdown of excitation-inhibition balance (Fig S4B (S1 Appendix)), establishing an increasingly excitatory system. Overall, Motif 2 experiences a decrease in inhibition with growing ionic imbalances. This mechanism has shown to be the precursor of epileptiform activities (55).

#### 3.1.3 Cellular ATP deficit

To simulate the effects of cellular ATP deficit, the parameter ATP was varied from 30μM to 0.01μM. The values 0.01μ to 15μM are varied in steps of 0.5μM and 15μM to 30μM in steps of 1μM. The Γ trends of neurons in both motifs is maintained at a steady value up to 5μM, after which the effect of lowering cellular ATP starts influencing the firing behavior. Hence, as seen in Fig 2C and Fig 2F, the phase *ATP* ≥ 5*μM* has been marked as *Normal phase*.

As the cellular ATP available decreases in Motif 1, at *ATP* ≤ 5*μM*, the Γ trends of both PYR1 and PYR2 increase as seen in Fig 2C and Fig S5B (S1 Appendix), marking the beginning of the *Critical phase*. Reduction in the available ATP causes the reduction in the activity of the active pumps, making it easier to depolarize. At 1.01μM, both the neurons exhibit synchronized firing rate trends, with Γ reaching a peak at 0.51μM (Fig S5A (S1 Appendix)). Neuronal activity thereby dangerously increases as the ATP deficit increases enforcing a detrimental positive feedback loop (24) . The neurons lose the ability to maintain resting membrane potential below firing threshold for *ATP* ≤ 0.51*μM* due to loss of ion gradients caused by active pump failure, leading to apoptotic cell death.

In motif 2, PYR is protected from the deleterious effects of hyperexcitation caused due to reduced cellular ATP by INH activity. From Fig 2F, in 2.01*μM* ≤ *ATP* ≤ 5*μM*, we observe a slight decrease in INH activity due to its sensitivity to growing ion imbalance. Nevertheless, INH imminently curtails the activity of PYR in this phase preventing excessive depolarizations (Fig S6C, (S1 Appendix)). In 1.01*μM* ≤ *ATP* ≤ 2.01*μM*, Γ of INH undergoes a massive spike due to reduced active pump functionality and loss of resting membrane potential.

The same spike is prevented in PYR because of increased INH activity (Fig S6B, (S1 Appendix)). 1.01*μM* ≤ *ATP* ≤ 5*μM*, has been marked as *Subcritical phase (self-restoring)* due to the ability of system to regulate the activity of PYR neuron. At points *ATP* ≤ 1.01*μM*, the activity of both the neurons ceases, marking the *Critical phase*. At 0.51*μM*, (Fig S6A, (S1 Appendix)), the INH activity is reduced to subthreshold (V<5mV) oscillations due to loss of ion gradients, but these oscillatory activities cause GABAergic activity, thereby causing the suppression of plausible hyperactivity in PYR (which is seen in Motif 1 for the same ATP value). Hence, INH activity substantially reduces the metabolic load in PYR during an energy crisis, provided other SBI processes do not reduce INH activity.

### 3.2 Restoration of firing behavior during ACS and DBS

ACS and DBS was applied at key events based on Γ trends of SBI processes as seen in Fig 2. The points of application of ACS and DBS has been marked in Fig 2.

#### 3.2.1 Motif 1

In motif 1, as shown in Fug. 1E, administration of ACS at certain points during SBI causes *reduction in Γ* as well as the *desynchronization* or *‘decoupling’* of PYR1 and PYR2 firing. Fig 3A shows Γ heatmap for various ACS amplitudes (y-axis) and ACS frequencies (x-axis) at [*Glu*^−^]*_e_* = 0.2*mM* for which Γ trends of PYR1 and PYR2 converge and synchronize. The inset figures Fig 3A (a_1_, a_2_), Fig 3A (b_1_, b_2_) show Γ at stim parameters like (20Hz, 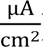 ), (20Hz, 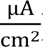 ), where there is a marginal decrease in Γ of both PYR1 and PYR2 from their pre-stimulation values (Fig S7A, (S1 Appendix)). At points such as (10Hz, 1.5 μA/cm^2^) and (120Hz, 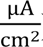), PYR2 fires at much lower value than PYR1, thereby *desynchronizing (decoupling)* the neurons (trace plot Fig S7B,C (S1 Appendix)). During this scenario, the phase of the system tends towards the *Normal phase*, with the PYR2 exhibiting near normal firing behavior. At 50Hz and 60Hz, both PYR1 and PYR2 exhibit very high ‘*resonant’* firing (Fig S7D, (S1 Appendix)) for high stimulation amplitude across different SBI processes and its various phases. Overall, such parameters, are detrimental for the system which increases the depolarizations instead of reducing them. At higher levels of [*Glu*^−^]*_e_*, PYR1 and PYR2 remain synchronized across ACS parameter space, with very few parameters being effective in curbing excessive depolarizations (In Fig S12A (S1 Appendix), for [*Glu*^−^]*_e_* = 0.45*mM*, only a small patch in firing rate heatmap shows neurons desynchronize with reduced Γ. At this point the motif firing behavior migrates from *Critical phase* to *Subcritical phase* behavior). ACS was applied at *ATP* = 0.51*μM*, a point in the *Critical phase* where both the PYR1 and PYR2 are synchronized and firing at very high Γ (Fig 2C). In Fig 3B, Γ for various ACS parameters at *ATP* = 0.51*μM* is shown. The inset figures Fig 3B(c_1_ , c_2_ ) show a set of parameters such as (15Hz, 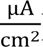 during which *excessive firing is reduced but not desynchronized* (Fig S8B, (S1 Appendix)). Here, the motif migrates closer to *Normal phase* firing behavior but there is lack of desynchronization of PYR1 and PYR2. Non-ideal parameter sets with low frequencies, moderate-high amplitudes such as (0.5Hz, 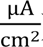 ) causes depolarization blocks due to reversal of ion gradient causing the system to stop functioning (Fig S8A, (S1 Appendix)). These manifest as patches of low firing activity seen in Fig 3B. Increasing *[K^+^]_o,rest_* causes an overall ionic imbalance in the system. It was observed that ACS or DBS paradigms proved relatively less effective for Motif 1 at high levels of *[K^+^]_o,rest_*, eliciting a minor change in the firing behavior of the system. As can be seen in Fig S12B (S1 Appendix), at 10mM of *[K^+^]_o,rest_*, application of various ACS parameters does not really change Γ. Further, the columns of resonant activity at 50Hz and 60Hz that are prominent in ATP and [*Glu*^−^]*_e_* firing rate responses for ACS, are significantly reduced.

**Fig 3:**
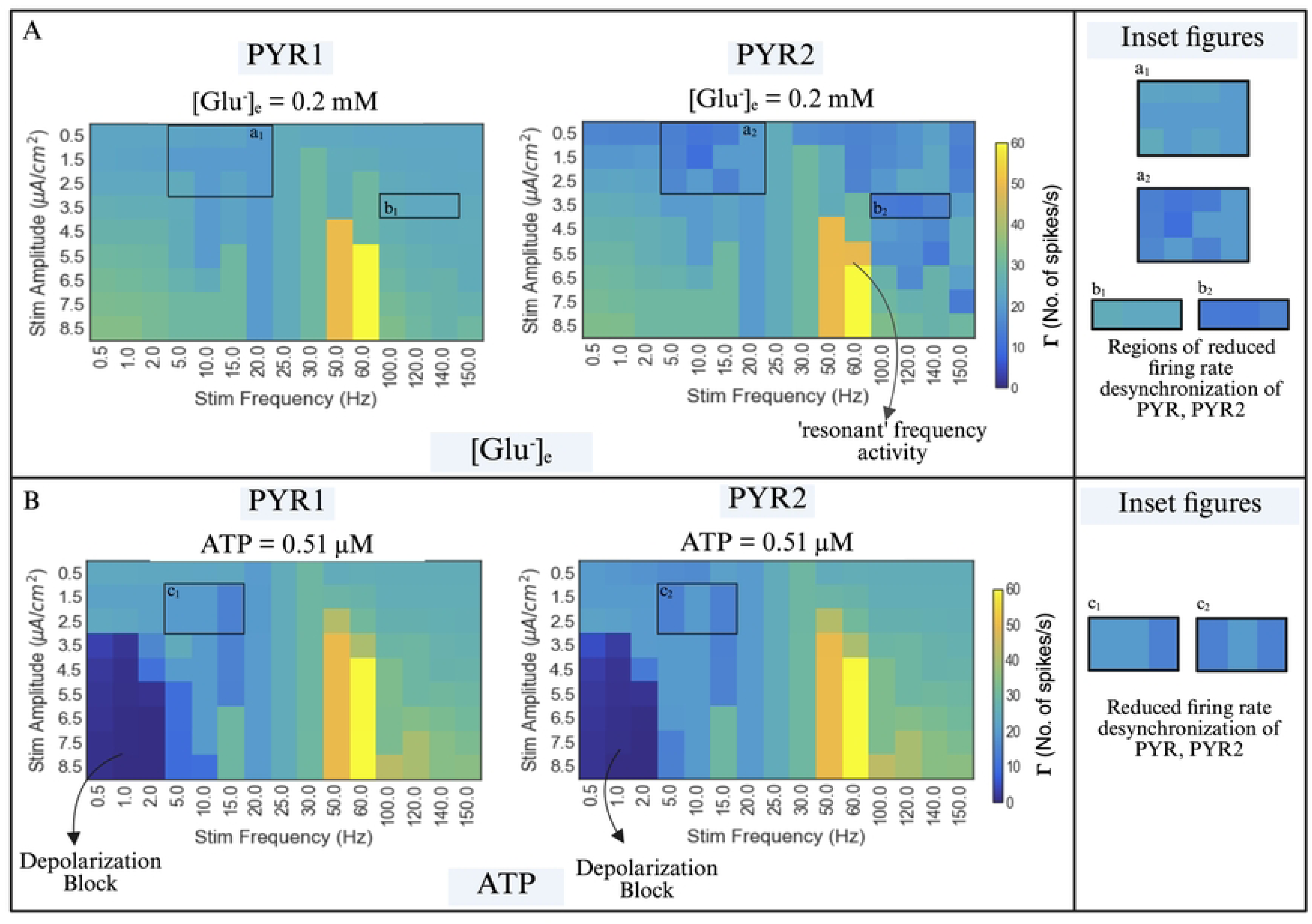
Firing rate responses for ACS parameters in Motif 1 at key SBI process events. **A**, The panel shows two heatmaps representing the firing rate responses for various ACS stimulation parameters (frequency on the x-axis and amplitude on the y-axis) at [*Glu*^−^]*_e_* = 0.2*mM*. The set of inset figures (a_1_, a_2_) showcase the ACS parameters at which reduced Γ and desynchronization of PYR1 and PYR2 occur and (b_1_,b_2_) showcases the points at which PYR2 desynchronizes from PYR1. The rest of the space is marked by increased depolarizations indicating worsening of conditions. **B,** The panel shows two heatmaps representing the firing rate responses for various ACS stimulation parameters at *ATP* = 0.51*μM*. At low frequency, moderate-high amplitude of ACS stimulation, we observe huge patches of Γ∼0Hz. These ‘pillars’ indicate low activity represent depolarization block. In this phase the system loses ionic gradient which disables transmission of action potentials. The set of inset figures (c_1_, c_2_) showcases ACS parameters at which Γ is reduced and desynchronized. The pillars of high activity at 50Hz, and 60Hz, triggered by moderate-high ACS amplitudes represents the *resonant* frequencies for the given neuronal model.

Monophasic DBS was similarly administered but due to nature of Motif 1, no stim parameters were proven to be useful on observing Γ. With PYR1 and PYR2 being pyramidal neurons interacting with glutamatergic synapse, monophasic DBS couples the system and increases Γ during SBI processes. In Fig 4, at *ATP* = 1.51*μM*, the system is not synchronized pre-stimulation but synchronizes instantly with much higher Γ trends across DBS amplitudes and frequencies. We notice similar trends in Fig S13A,B ([*Glu*^−^]*_e_*, [*K*^+^]_*o,rest*_) (S1 Appendix) where PYR1 and PYR2 exhibit synchronized firing with exceeding DBS amplitudes.

**Fig 4:**
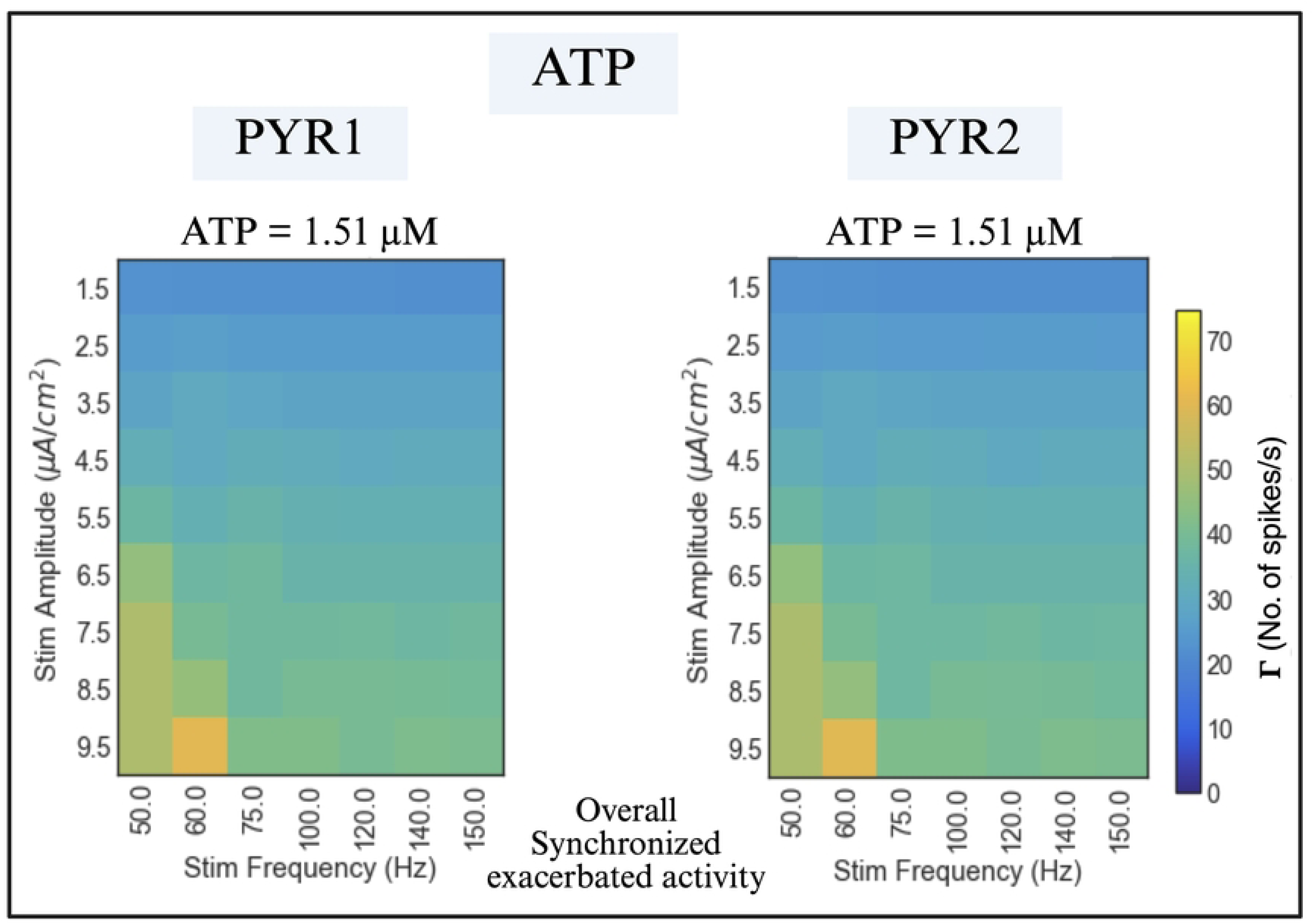
The effect of DBS parameters at *ATP* = 1.5*μM* in Motif 1. We see here that PYR1 and PYR2 synchronize in their activities early on along DBS amplitude values.

#### 3.2.2 Motif 2

Along with implementing various combinations of amplitudes and frequencies, 3 different stimulation strategies were implemented on Motif 2 due to the different categories of neurons present (1 pyramidal, 1 interneuron) as seen in Fig 1E. The 3 strategies are: Stimulating *PYR-INH together*, stimulating *only PYR*, stimulating *only INH*. These strategies were implemented for both ACS and DBS paradigms. It was observed that stimulating only PYR worsened the firing behavior of PYR from the pre-stimulation condition during SBI.

Fig 5A shows [*Glu*^−^]*_e_* = 0.29*mM* under PYR-INH stim strategy. In the inset figures Fig 5A(a_1_, a_2_) we see reduced PYR activity with the system migrating closer to the *Subcritical phase (self-restoring)*. We observe huge ‘pillars’ of Γ∼0 at 25Hz, 30Hz, 50Hz and 60Hz in INH firing heatmap. At these stim frequencies and amplitudes, depolarizations of INH increases dramatically eventually reducing INH firing activity to mere sub-threshold oscillations (V<5mV), which continues to induce GABAergic activity. This causes strong suppression of PYR firing (Fig S9, (S1 Appendix)), thereby subduing spurious depolarizations induced by [*Glu*^−^]*_e_*. The ACS INH strategy was found to elicit better firing rate responses at [*K*^+^]_*o,rest*_ = 5.8*mM* than PYR-INH strategy (where, firing rate of PYR worsens). In the inset figures Fig 5(b_1_, b_2_), Fig 5(c_1_, c_2_) we see an increase in INH activity which suppresses PYR activity marginally. During ATP deficit, INH activity suppresses the excessive depolarizations of PYR thereby preventing hyperactivity. Thus, in such self-restoring subcritical situations, electrical stimulation of any kind is not necessary. Fig S14A (S1 Appendix) showcases the effects of ACS paradigms under INH stim strategy at *ATP* = 1.51*μM*. It further reduces PYR activity suppression at 30Hz, 50Hz, 60Hz due excessive INH stimulation and subthreshold oscillation induced GABAergic activity.

**Fig 5:**
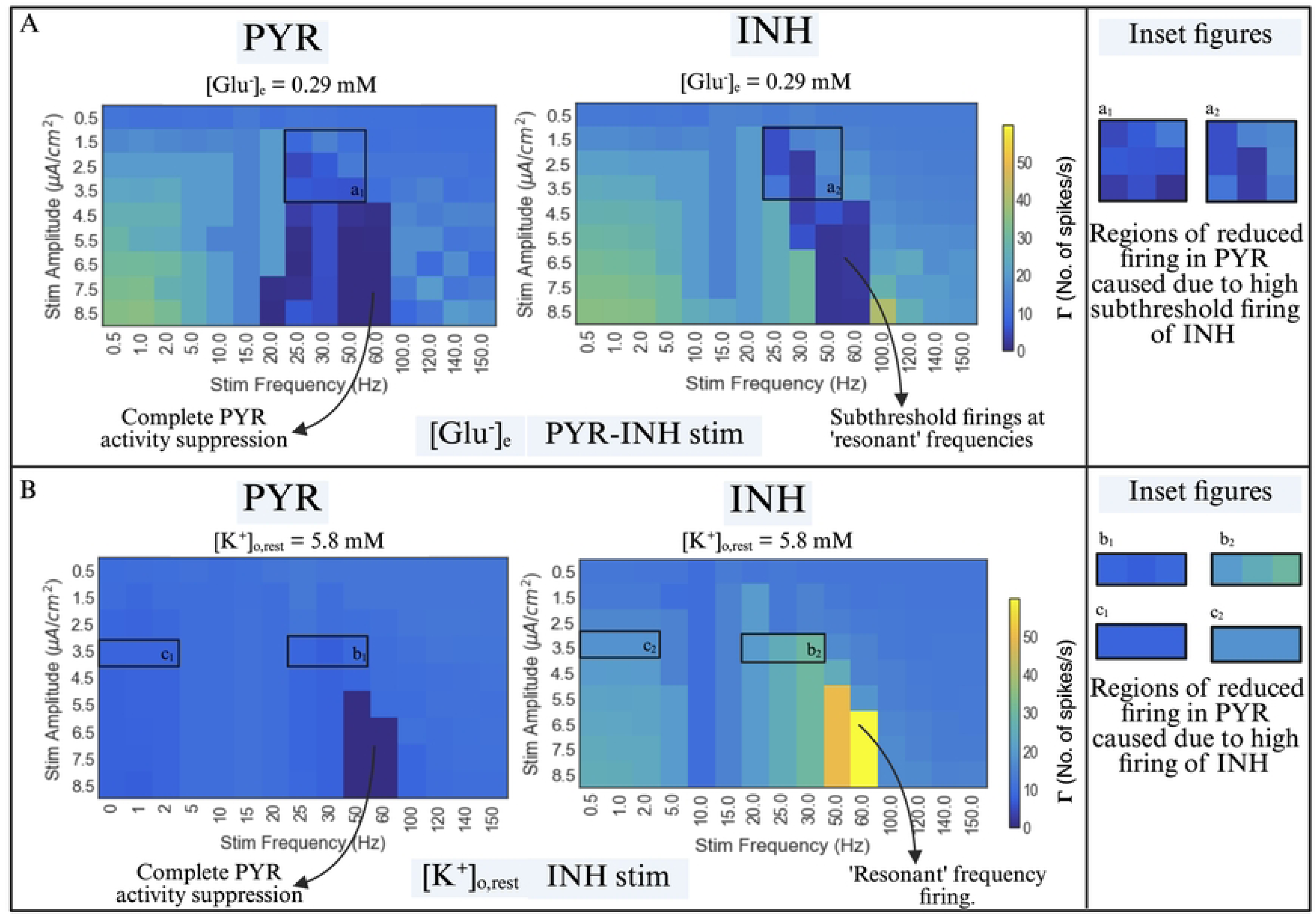
The firing rate responses for different ACS parameters and strategies in Motif 2 at key SBI events. The best result in each strategy is presented. **A,** The firing rate response heatmaps shows the effect of ACS paradigms in PYR-INH stim strategy at [*Glu*^−^]*_e_* = 0.29*mM*. The set of inset figures (a_1_, a_2_) highlights the increased activity in INH neuron which causes subthreshold oscillations in INH which induces GABAergic activity thereby restoring the PYR firing nearly to its normal value. **B,** The firing rate response heatmaps show the effect of ACS paradigms in INH strategy at [*K*^+^]_*o,rest*_ = 5.8*mM* in Motif 2. The set of inset figures (b_1_, b_2_) and (c_1_, c_2_) show increased INH activity which moderately reduces PYR activity. Increase in [*K*^+^]_*o,rest*_ makes the system a bit resistant to the effects of stimulation. Here, we see that certain parameters easily cause INH activity, but complete PYR activity suppression only occurs at the resonant frequencies where the INH activity is primarily increased. We see marginal separation in other areas shown in the inset figures.

For monophasic DBS, it was observed that the strategy stimulating only INH gave the best results. In Fig 6A, at [*Glu*^−^]*_e_* = 0.29*mM*, we see uniformly decreasing Γ for increasing amplitude across various frequencies of DBS. In the inset figures Fig 6A(a_1_, a_2_), we see sustained suppression of PYR activity due to increased INH activity and subthreshold oscillations, wherein the firing activity of PYR has almost returned to *Normal phase*. Fig S10 (S1 Appendix) shows the action potential traces showing this effect at (100Hz, 4.5 μA/cm^2^). For ATP deficit, we see that monophasic DBS paradigms with INH strategy proves useful albeit not a necessity. Inset figures Fig 6B(b_1_, b_2_) show that the increased INH activity due to DBS causes sustained PYR activity suppression. The action potential traces at (60Hz, 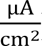 ) (Fig S11 (S1 Appendix)) shows the same. This prevents downstream propagation of sustained PYR activity and reduces the overall metabolic load. The dark patches with low Γ at higher DBS amplitudes for INH in Fig 6A,B indicate subthreshold oscillations which is caused due to high DBS frequencies and amplitudes. Activities of such nature cannot be considered as conventional firing, but does lead to GABAergic activity, thereby suppressing PYR activity completely as seen in Fig 6. Fig S14B (S1 Appendix) shows the effects of DBS at [*K*^+^]_*o,rest*_ = 5.8*mM*. With increasing DBS amplitudes, Γ increases for INH, but it does not affect the Γ of PYR drastically. The firing rate response space of PYR for DBS is unvariegated except for some drastic activity suppressions at high amplitudes. This highlights the effects of [*K*^+^]_*o,rest*_ on the motif, making it resilient to external stimuli.

**Fig 6:**
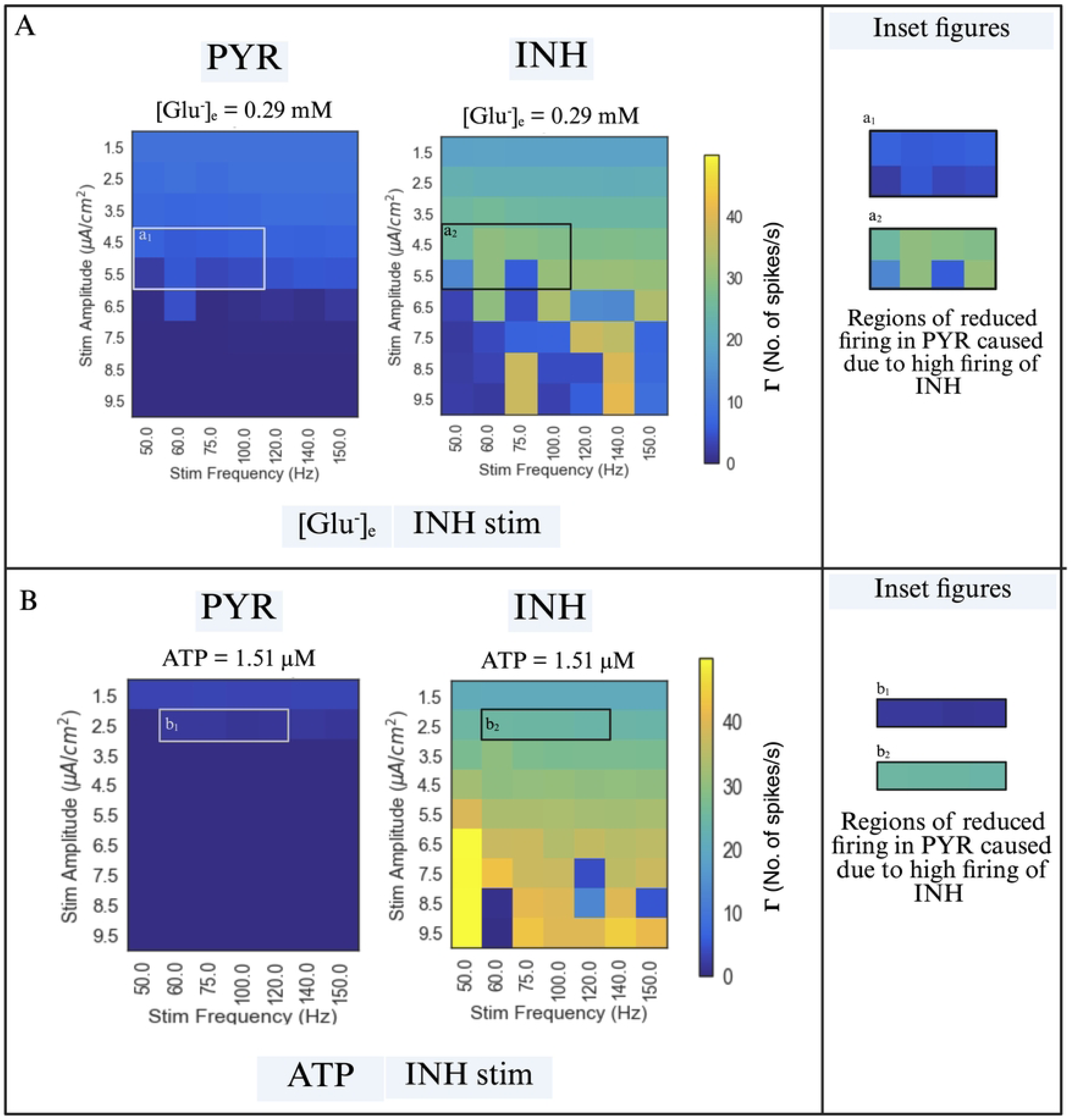
The firing rate responses for DBS parameters and strategies in Motif 2 at key SBI events. INH stim strategy works the best with DBS paradigms. **A,** The figure shows the firing rate responses of Motif 2 at [*Glu*^−^]*_e_* = 0.29*mM*. In the inset figure (a_1_, a_2_) we see that INH stim increases INH Γ, exhibiting subthreshold activations at 50Hz, 75Hz. This in turn suppresses PYR activity and thereby spurious downstream activity. **B,** The figure shows the firing rate responses of Motif 2 at *ATP* = 1.51*μM*. Although pre stimulation INH activity is high enough to control PYR activity, we see here that DBS further stimulates INH neuron causing high suppression of PYR activity at lower DBS stimulation amplitudes.

## 4. Mechanisms of stimulation during firing behavior restoration

### 4.1 ACS vs. DBS

ACS induces *direct sustained changes* in *V_m_* (membrane potential of the neurons). This leads to a variegated stimulation parameter space (Fig 3, Fig S12 (S1 Appendix)). This mechanism of ACS causes *desynchronization* of neurons in Motif 1 (Fig S7B,C (S1 Appendix)). The appropriate ACS parameters suppress high frequency neural firings in Motif 1 during different phases of SBI, and the inappropriate ACS parameters exacerbate Γ leading to excessive firing discharges resembling non-epileptiform activity (Fig S7D (S1 Appendix)). The ACS amplitudes at which Γ improves or worsens depends on the phase of SBI process dominating the system.

Further, across the ACS parameter space, the frequency parameter is the dominant variable (Fig 3,5, Fig S12 (S1 Appendix)). Frequencies like 5Hz, 10Hz, 15Hz, 20Hz, 25Hz, 30Hz often restore Γ for Motif 1 across SBI parameters thereby changing the phase of the system. These frequencies are commonly used in experimental ACS paradigms. Certain high frequency parameters such as 120Hz, do show similar improvements in some cases (Fig 3A). Moreover, the firing behavior of both motifs during ACS, show a *model-specific* nature: changing the Hodgkin Huxley formulation by altering ion channel category and dynamics could change the firing rate response of the system with the ‘resonant’ frequencies and other important ACS parameters being altered. In Motif 2, it was observed that INH stim strategy with ACS worked the best for early phases of SBI and PYR-INH stim strategy with ACS worked the best for advanced phases of SBI. This can be attributed to the nature of ACS waveform and the phase of the SBI. Overall, ACS is more versatile because there is a chance of finding the best set of parameters (the set of parameters for which the PYR exhibit a normal Γ or a reduction in hyperactivity, thereby altering the phase of the motif from *Subcritical/Critical* to *Normal/Subcritical*) for stimulation at a given stage of SBI process. The disadvantage here is the parameters adjacent to the good set of parameters space could be detrimental to the motif by exacerbating the excitability.

Monophasic DBS waveform consists of very narrow pulses of charges (electrical current) in the positive phase, injected into neural tissues which induce a transient change in the membrane potential instead of sustained change as in ACS. Due to this, DBS waveform easily excites and synchronizes a feedforward excitatory system like Motif 1 (Fig 4). Hence, administration of Monophasic DBS is not useful in a system dominated by excitatory neurons and synapses. It was found to act through synaptic transmission rather than sustained modulation of the membrane potential as in the case of ACS. In Motif 2, stimulation using the INH stim strategy proves to be the best for all the cases because it acts through INH, thereby inducing increased GABAergic transmission in the system. This makes it effective during *Critical phases* of the SBI processes leading to PYR hyperactivity suppression. Stimulation using PYR-INH strategy exacerbates the PYR activity under normal ranges of amplitude and frequency parameters. Firing rate response in the Monophasic DBS parameter spaces is more uniform than ACS (Fig 4 ,Fig 6). The frequencies tested are the standard frequencies used in experiments, hence most of the frequencies show promising response. Overall, effects of monophasic DBS in the local network of neurons depends on the target neuron category and the underlying nature of the local synaptic transmission, i.e., whether the target is interneuron with local inhibitory activity or principal neurons with local excitatory activity.

ACS is suitable for stimulating generically when the underlying properties of motifs such as type of neurons, nature of synapses and connectivity is not well known. Though the firing rate response space for ACS parameters is dynamic, there is a reasonable chance of opting decent parameters with each trial. Monophasic DBS works best when the properties of target neurons and local synaptic transmission is known and is easier to modify the choice of stimulation parameters without extreme changes in firing rate responses.

### 4.2 Motif 1 vs. Motif 2

Motif 1, being a solely feedforward excitatory system is unable to control or restore the firing rate response of its own neurons during various stages of the SBI processes due to lack of internal control. As can be seen in Fig 2A,B,C, PYR1 and PYR2 exhibit increased firing activity as SBI progresses, synchronizing PYR1-PYR2.

Modulating this neural activity, desynchronizing PYR1, PYR2, though achieved for certain parameters by ACS, is a difficult task. Without external intervention, sustained depolarizations eventually lead to loss of ion gradients, intracellular calcium ion accumulation, and cellular death. Though such motifs seem irredeemable, we show that ACS parameter space exhibits stim parameters that curb sustained depolarizations, thereby aiding it to change its phase even though conditions are not ideal. Realistically, this prevents PYR neurons in Motif 1 from entering neurodegenerative stages and reduces the metabolic load.

In Motif 2, due to feedforward GABAergic synapse induced by INH activity, the tendency of PYR to undergo excessive depolarizations during various phases of the SBI process is curbed. Due to this, during the initial stages of the SBI process, we see active suppression of spurious activity in PYR. This self-regulation makes the system resilient to initial stages of SBI. Worsening of SBI reduces inhibition due to reduced GABAergic activity, which is sensitive to changing ionic environment, thereby disrupting the *inherent excitation-inhibition balance*. By choosing suitable strategies (PYR-INH stim, or INH stim) of ACS or DBS, GABAergic transmission can be restored by increasing the INH activity. Though ACS is seen to work well, DBS shows immense promise in the INH stim strategy across a wide range of stimulation parameters.

Inhibition is a crucial element that is jeopardized during SBI. In fact, a solely feedforward excitatory neural motif exhibits the ideal conditions needed for the system to display non-epileptiform discharges during Critical phase. Ideally, increasing inhibitory activity during SBI processes, which is possible in systems like Motif 2, provided the inhibitory neuron has not completely lost its functionality, helps in countering consequences of SBI. Overall, Motif 2 responds more reliably to stimulation than Motif 1 during suitable SBI phases due to the presence of inhibitory element.

### 4.3 SBI processes and stimulation paradigms

The nature of the simulated SBI processes primarily involve increasing depolarizations in neurons. Hence, Γ was the most intuitive measure to understand the changes in motif behavior. During glutamate excitotoxicity, we show that the right ACS stim parameters suppresses excessive firing and desynchronizes PYR1 and PYR2 in Motif 1. But certain stimulation parameters could exacerbate hyperactivity during such conditions. During ATP deficit, bad stimulation parameters cause further excitability in Motif 1 leading to depolarization block. Further, ACS produces subthreshold oscillations during glutamate excitotoxicity in INH of Motif 2. Though it produces GABAergic inhibition during this process, sustained INH activity of such nature is undesirable. ATP was found to be the most severe SBI parameter, in terms of Γ trends in both motifs, but certain stimulation parameters show promise in restoring Γ to normalcy thereby reducing the metabolic load. On the other hand, [*K*^+^]_*o,rest*_, severely affected the ability of the motifs to respond to either ACS or DBS stimulation paradigms. At moderately high levels of [*K*^+^]_*o,rest*_, certain ACS and DBS parameters partially curb PYR hyperactivity, but the rest of the parameter space shows that electrical stimulation exacerbates the situation. Further, high levels of [*K*^+^]_*o,rest*_caused the motifs to be less responsive to stimulation, generating a smoother parameter space with sustained high depolarizations. Overall, phase of SBI process seemed to affect ACS paradigms more than DBS. Choosing the right stimulation strategy for DBS modulated PYR response despite being in *Subcritical/Critical* phases.

Our work has delineated the effects of the SBI processes in two neuron motifs. The feedforward excitatory and inhibitory motifs are the building blocks of neuronal populations. Understanding SBI processes in fundamental motifs and the effects of electrical stimulation during such scenarios has not been studied. It is very intuitive to think that electrical stimulation further sensitizes the neural tissue by exacerbating depolarizations during SBIs because of the nature of the processes involved. However, there are specific stimulation parameters and strategies that produce an efficacious response and restore spiking activity to a healthy range. Furthermore, there are phases along SBI, where administration of ACS or DBS is effective. The neurons were not modeled to capture fatigue due to stimulation, the downstream effects in the glial-vascular units (56) or the complex intracellular pathways that are influenced during brain stimulations (57). It should also be noted the SBI processes modeled are often accompanied by other processes such as neuroinflammation, cytotoxic edema, oxidative stress, modifications in synaptic plasticity, which will be explored in future works. Overall, ACS and DBS have different mechanisms of action and are beneficial when applied during specific key events that differ between the stimulation types. Though this work discusses the effects of single SBI processes, trends of firing rates for pairs of SBI processes were also simulated. When multiple SBI processes are superimposed, points along firing rate trends at which ACS or DBS could restore firing behavior decreased. This is shown to be true in real scenarios wherein multiple SBI processes occur in tandem, making it challenging to find a suitable opportunity to administer stimulation. Nevertheless, this work offers a proof of concept that the evolution of the SBI phases in both the motifs, does provide us with some opportune moments to administer suitable stimulation paradigms that exhibit promising prognosis.

## Acknowledgements

We thank Houston Methodist Hospital and Houston Methodist Academic Institute for supporting and funding this work. All the images were drafted using Biorender (https://BioRender.com).

## Supporting Information caption

S1 Appendix. The supporting information file consists of detailed methods, equations, tables of constants and additional figures for providing further clarity of results.

## Data availability statement

### Code reporting

All code written in support of this publication is publicly available in a Zenodo repo (https://doi.org/10.5281/zenodo.17955177) and all data generated, and images plotted are available in Figshare (10.6084/m9.figshare.30896399). Any additional information required to reanalyze the data reported in this paper is available from the lead contact upon request.

## References

1. Zhong H, Feng Y, Shen J, Rao T, Dai H, Zhong W, et al. Global Burden of Traumatic Brain Injury in 204 Countries and Territories From 1990 to 2021. Am J Prev Med [Internet]. 2025;68(4):754–63. Available from: https://www.sciencedirect.com/science/article/pii/S0749379725000017

2. Feigin Valery L, Brainin Michael, Norrving Bo, Martins Sheila O, Pandian Jeyaraj, Lindsay Patrice, et al. World Stroke Organization: Global Stroke Fact Sheet 2025. International Journal of Stroke [Internet]. 2025 Jan 3;20(2):132–44. Available from: 10.1177/17474930241308142

3. Shao L, Chen S, Ma L. Secondary Brain Injury by Oxidative Stress After Cerebral Hemorrhage: Recent Advances. Front Cell Neurosci [Internet]. 2022;Volume 16-2022. Available from: https://www.frontiersin.org/journals/cellular-neuroscience/articles/10.3389/fncel.2022.853589

4. Rana A, Singh S, Sharma R, Kumar A. Traumatic Brain Injury Altered Normal Brain Signaling Pathways: Implications for Novel Therapeutics Approaches. Curr Neuropharmacol [Internet]. 2019;17:614–29. Available from: https://api.semanticscholar.org/CorpusID:52189835

5. Bautista Wendy, Adelson P. David, Bicher Nathan, Themistocleous Marios, Tsivgoulis Georgios, Chang Jason J. Secondary mechanisms of injury and viable pathophysiological targets in intracerebral hemorrhage. Ther Adv Neurol Disord [Internet]. 2021 Jan 1;14:17562864211049208. Available from: 10.1177/17562864211049208

6. Duris K, Splichal Z, Jurajda M. The role of inflammatory response in stroke associated programmed cell death. Curr Neuropharmacol. 2018;16(9):1365–74.

7. Chesnut RM, Marshall LF, Klauber MR, Blunt BA, Baldwin N, Eisenberg HM, et al. THE ROLE OF SECONDARY BRAIN INJURY IN DETERMINING OUTCOME FROM SEVERE HEAD INJURY. Journal of Trauma and Acute Care Surgery [Internet]. 1993;34(2). Available from: https://journals.lww.com/jtrauma/fulltext/1993/02000/the_role_of_secondary_brain_injury_in_determining.6.aspx

8. Datusalia A, Khatri N, Kaundal R, Sharma S, Kumar DS. The Complexity of Secondary Cascade Consequent to Traumatic Brain Injury: Pathobiology and Potential Treatments. Curr Neuropharmacol. 2019 Dec 12;19.

9. Seule M, Brunner T, Mack A, Hildebrandt G, Fournier JY. Neurosurgical and intensive care management of traumatic brain injury. Facial plastic surgery. 2015;31(04):325–31.

10. Mustafa AG, Alshboul OA. Pathophysiology of traumatic brain injury. Neurosciences Journal. 2013;18(3):222–35.

11. Beez T, Steiger HJ, Etminan N. Pharmacological targeting of secondary brain damage following ischemic or hemorrhagic stroke, traumatic brain injury, and bacterial meningitis - a systematic review and meta-analysis. BMC Neurol [Internet]. 2017;17(1):209. Available from: 10.1186/s12883-017-0994-z

12. Schiff ND, Giacino JT, Butson CR, Choi EY, Baker JL, O’Sullivan KP, et al. Thalamic deep brain stimulation in traumatic brain injury: a phase 1, randomized feasibility study. Nat Med [Internet]. 2023;29(12):3162–74. Available from: 10.1038/s41591-023-02638-4

13. Cordeiro BN de L, Rocha JVDS, Kuster E, Thibauth A, Nascimento LR, Swank C, et al. Transcranial direct current stimulation in individuals with severe traumatic brain injury in the subacute phase: a case series. Front Hum Neurosci [Internet]. 2025;Volume 19-2025. Available from: https://www.frontiersin.org/journals/human-neuroscience/articles/10.3389/fnhum.2025.1552387

14. Ziesel D, Nowakowska M, Scheruebel S, Kornmueller K, Schäfer U, Schindl R, et al. Electrical stimulation methods and protocols for the treatment of traumatic brain injury: a critical review of preclinical research. J Neuroeng Rehabil [Internet]. 2023;20(1):51. Available from: 10.1186/s12984-023-01159-y

15. Cha S, Choi J, Moon C, Cho K. Non-invasive brain stimulation contributing to postural control with and without stroke: a systematic review and meta-analysis. Sci Rep [Internet]. 2025;15(1):26020. Available from: 10.1038/s41598-025-07840-7

16. Li KP, Wu JJ, Zhou ZL, Xu DS, Zheng MX, Hua XY, et al. Noninvasive Brain Stimulation for Neurorehabilitation in Post-Stroke Patients. Brain Sci [Internet]. 2023;13(3). Available from: https://www.mdpi.com/2076-3425/13/3/451

17. Hofmeijer J, Ham F, Kwakkel G. Evidence of rTMS for Motor or Cognitive Stroke Recovery: Hype or Hope? Stroke [Internet]. 2023 Oct 1;54(10):2500–11. Available from: 10.1161/STROKEAHA.123.043159

18. Baker KB, Plow EB, Nagel S, Rosenfeldt AB, Gopalakrishnan R, Clark C, et al. Cerebellar deep brain stimulation for chronic post-stroke motor rehabilitation: a phase I trial. Nat Med [Internet]. 2023;29(9):2366–74. Available from: 10.1038/s41591-023-02507-0

19. Malakouti N, Serruya MD, Cramer SC, Kimberley TJ, Rosenwasser RH. Making Sense of Vagus Nerve Stimulation for Stroke. Stroke [Internet]. 2024 Feb 1;55(2):519–22. Available from: 10.1161/STROKEAHA.123.044576

20. Hodgkin AL, Huxley AF. A quantitative description of membrane current and its application to conduction and excitation in nerve. J Physiol. 1952;500–44.

21. Wei Y, Ullah G, Schiff SJ. Unification of Neuronal Spikes, Seizures, and Spreading Depression. Journal of Neuroscience [Internet]. 2014;34(35):11733–43. Available from: https://www.jneurosci.org/content/34/35/11733

22. Song J, Kim J, Struck A, Zhang R, Westover MB. A Model of Metabolic Supply-Demand Mismatch Leading to Secondary Brain Injury. J Neurophysiol. 2021 Jul 7;126.

23. Cressman JR, Ullah G, Ziburkus J, Schiff SJ, Barreto E. The influence of sodium and potassium dynamics on excitability, seizures, and the stability of persistent states: I. Single neuron dynamics. J Comput Neurosci [Internet]. 2009;26(2):159–70. Available from: 10.1007/s10827-008-0132-4

24. Le Masson G, Przedborski S, Abbott LF. A Computational Model of Motor Neuron Degeneration. Neuron [Internet]. 2014;83(4):975–88. Available from: https://www.sciencedirect.com/science/article/pii/S0896627314005820

25. Aberra AS, Wang B, Grill WM, Peterchev A V. Simulation of transcranial magnetic stimulation in head model with morphologically-realistic cortical neurons. Brain Stimul [Internet]. 2020;13(1):175–89. Available from: https://www.sciencedirect.com/science/article/pii/S1935861X19304097

26. Mirzakhalili E, Barra B, Capogrosso M, Lempka SF. Biophysics of Temporal Interference Stimulation. Cell Syst [Internet]. 2020;11(6):557–572.e5. Available from: https://www.sciencedirect.com/science/article/pii/S2405471220303720

27. Ahsan F, Govindaraju AC, Raphael RM, Chi T, Sheth SA, Goodman W, et al. Biophysics of amplitude-modulated gigahertz electromagnetic waves stimulation. In: 2023 57th Asilomar Conference on Signals, Systems, and Computers. 2023. p. 1463–8.

28. Rubin JE, Terman D. High frequency stimulation of the subthalamic nucleus eliminates pathological thalamic rhythmicity in a computational model. J Comput Neurosci [Internet]. 2004;16(3):211–35. Available from: http://europepmc.org/abstract/MED/15114047

29. Muddapu VR, Mandali A, Chakravarthy VS, Ramaswamy S. A Computational Model of Loss of Dopaminergic Cells in Parkinson’s Disease Due to Glutamate-Induced Excitotoxicity. Front Neural Circuits [Internet]. 2019;Volume 13-2019. Available from: https://www.frontiersin.org/journals/neural-circuits/articles/10.3389/fncir.2019.00011

30. Izhikevich EM. Simple model of spiking neurons. IEEE Trans Neural Netw. 2003;14(6):1569–72.

31. Khodashenas M, Baghdadi G, Towhidkhah F. A modified Hodgkin–Huxley model to show the effect of motor cortex stimulation on the trigeminal neuralgia network. The Journal of Mathematical Neuroscience [Internet]. 2019;9(1):4. Available from: 10.1186/s13408-019-0072-5

32. Zhou J, Khateeb K, Yazdan-Shahmorad A. Early intervention with electrical stimulation reduces neural damage after stroke in non-human primates. Nat Commun [Internet]. 2025;16(1):6701. Available from: 10.1038/s41467-025-61948-y

33. Byrne JH, Heidelberger R, Waxham MN. From molecules to networks: an introduction to cellular and molecular neuroscience. Academic Press; 2014.

34. Tegner J, Compte A, Wang XJ. The dynamical stability of reverberatory circuits. Biol Cybern. 2003 Aug;87:471–81.

35. Wang XJ, Tegné J, Constantinidis C, Goldman-Rakic PS, Raichle ME. Division of labor among distinct subtypes of inhibitory neurons in a cortical microcircuit of working memory [Internet]. Vol. 101. 2004. Available from: www.pnas.orgcgidoi10.1073pnas.0305337101

36. Chander Bankim Subhash AND Chakravarthy VS. A Computational Model of Neuro-Glio-Vascular Loop Interactions. PLoS One [Internet]. 2012 Aug;7(11):1–11. Available from: 10.1371/journal.pone.0048802

37. Destexhe A, Mainen Z, Sejnowski T. Kinetic Models of Synaptic Transmission. Vol. 2, Methods in Neuronal Modelling, From Ions to Networks. 1998. 1–26 p.

38. Magi S, Piccirillo S, Amoroso S, Lariccia V. Excitatory Amino Acid Transporters (EAATs): Glutamate Transport and Beyond. Int J Mol Sci [Internet]. 2019;20(22). Available from: https://www.mdpi.com/1422-0067/20/22/5674

39. Flanagan B, McDaid L, Wade J, Wong-Lin K, Harkin J. A computational study of astrocytic glutamate influence on post-synaptic neuronal excitability. PLoS Comput Biol [Internet]. 2018 Apr 16;14(4):e1006040-. Available from: 10.1371/journal.pcbi.1006040

40. Katayama Y, Becker DP, Tamura T, Hovda DA. Massive increases in extracellular potassium and the indiscriminate release of glutamate following concussive brain injury. J Neurosurg [Internet]. 1990;73(6):889–900. Available from: https://thejns.org/view/journals/j-neurosurg/73/6/article-p889.xml

41. Reinert M, Khaldi A, Zauner A, Doppenberg E, Choi S, Bullock R. High extracellular potassium and its correlates after severe head injury: relationship to high intracranial pressure. Neurosurgical Focus FOC [Internet]. 2000;8(1):1–8. Available from: https://thejns.org/focus/view/journals/neurosurg-focus/8/1/foc.2000.8.1.2027.xml

42. Zhang X, Zhang Y, Su Q, Liu Y, Li Z, Yong VW, et al. Ion Channel Dysregulation Following Intracerebral Hemorrhage. Neurosci Bull [Internet]. 2024;40(3):401–14. Available from: 10.1007/s12264-023-01118-6

43. Sick Thomas J, Feng Zi-Cai, Rosenthal Myron. Spatial Stability of Extracellular Potassium Ion and Blood Flow Distribution in Rat Cerebral Cortex after Permanent Middle Cerebral Artery Occlusion. Journal of Cerebral Blood Flow & Metabolism [Internet]. 1998 Oct 1;18(10):1114–20. Available from: 10.1097/00004647-199810000-00008

44. Chang RCC, Hudson PM, Wilson BC, Liu B, Abel H, Hong JS. High concentrations of extracellular potassium enhance bacterial endotoxin lipopolysaccharide-induced neurotoxicity in glia–neuron mixed cultures. Neuroscience [Internet]. 2000;97(4):757–64. Available from: https://www.sciencedirect.com/science/article/pii/S0306452200000592

45. Badawe HM, El Hassan RH, Khraiche ML. Modeling ultrasound modulation of neural function in a single cell. Heliyon [Internet]. 2023;9(12):e22522. Available from: https://www.sciencedirect.com/science/article/pii/S240584402309730X

46. Clewley R. Hybrid Models and Biological Model Reduction with PyDSTool. PLoS Comput Biol [Internet]. 2012 Aug 9;8(8):e1002628-. Available from: 10.1371/journal.pcbi.1002628

47. Hairer E, Wanner G. Solving Ordinary Differential Equations II. Stiff and Differential-Algebraic Problems. Vol. 14, Springer Verlag Series in Comput. Math. 1996.

48. Ankarcrona M, Dypbukt JM, Bonfoco E, Zhivotovsky B, Orrenius S, Lipton SA, et al. Glutamate-induced neuronal death: a succession of necrosis or apoptosis depending on mitochondrial function. Neuron. 1995;15(4):961–73.

49. Garcia JD, Gookin SE, Crosby KC, Schwartz SL, Tiemeier E, Kennedy MJ, et al. Stepwise disassembly of GABAergic synapses during pathogenic excitotoxicity. Cell Rep [Internet]. 2021;37(12):110142. Available from: https://www.sciencedirect.com/science/article/pii/S2211124721016387

50. Guerriero RM, Giza CC, Rotenberg A. Glutamate and GABA Imbalance Following Traumatic Brain Injury. Curr Neurol Neurosci Rep [Internet]. 2015;15(5):27. Available from: 10.1007/s11910-015-0545-1

51. Fertziger AP, Ranck Jr JB. Potassium accumulation in interstitial space during epileptiform seizures. Exp Neurol. 1970;26(3):571–85.

52. Florence G, Dahlem MA, Almeida ACG, Bassani JWM, Kurths J. The role of extracellular potassium dynamics in the different stages of ictal bursting and spreading depression: A computational study. J Theor Biol [Internet]. 2009;258(2):219–28. Available from: https://www.sciencedirect.com/science/article/pii/S0022519309000502

53. Hansen AJ, Zeuthen T. Extracellular ion concentrations during spreading depression and ischemia in the rat brain cortex. Acta Physiol Scand. 1981;113(4):437–45.

54. Wu X, Lu X, Lu X, Yu J, Sun Y, Du Z, et al. Prevalence of severe hypokalaemia in patients with traumatic brain injury. Injury [Internet]. 2015 Jan 1;46(1):35–41. Available from: 10.1016/j.injury.2014.08.002

55. Symonds C. Hughlings Jackson Lecture: Excitation and Inhibition in Epilepsy. SAGE Publications; 1959.

56. Muguli Ananya. The Biophysics of Neuron-Astrocyte-Vascular Modeling in Conditions of Normalcy, Intracerebral Hemorrhagic (ICH) Stroke and Electrical Stimulation. 2024;

57. Chen Y, Zhu G, Yuan T, Ma R, Zhang X, Meng F, et al. Subthalamic nucleus deep brain stimulation alleviates oxidative stress via mitophagy in Parkinson’s disease. NPJ Parkinsons Dis [Internet]. 2024;10(1):52. Available from: 10.1038/s41531-024-00668-4

